# Longitudinal analysis of the gut microbiome in the 5xfAD mouse model of Alzheimer’s disease

**DOI:** 10.1101/2022.03.02.482725

**Authors:** Sage J. B. Dunham, Katelyn A. McNair, Eric D. Adams, Julio Avelar-Barragan, Stefania Forner, Mark Mapstone, Katrine L. Whiteson

**Affiliations:** Department of Molecular Biology & Biochemistry University of California Irvine 3315 McGaugh Hall Irvine, CA, 92697; Department of Computational Science University of California Irvine 3019 Donald Bren Hall Irvine, CA, 92697; Institute for Memory Impairments and Neurological Disorders (UCI MIND) University of California Irvine Biological Sciences III, 2642 Irvine, CA, 92697; Department of Neurology University of California Irvine Irvine, CA, United States

**Keywords:** Alzheimer’s Disease; MODEL-AD, Microbiome, gut-brain axis, *Turicibacter*, Serotonin

## Abstract

**INTRODUCTION:** Microbial exposures impact Alzheimer’s disease (AD), and a baseline understanding of AD model microbiomes is vital for improving model efficacy. We describe the evaluation of the microbiome and metabolome in longitudinal samples from the 5xfAD transgenic mouse model.

**METHODS:** Cecal and fecal samples from 5xfAD and wild-type B6J (WT) animals from 4– 18 months of age were subjected to DNA extraction and shotgun Illumina sequencing. Metabolomics was performed on plasma and feces from a subset of animals.

**RESULTS:** Significant sex, age, and cage-specific differences were observed in the microbiome. Eight bacteria species were elevated in older 5xfAD mice and nine were depleted, including *Turicibacter* spp. Contradicting published findings covering persons with AD, plasma measurements revealed elevated serotonin in 18 month 5xfAD animals.

**DISCUSSION:** Taken together, these findings strengthen the link between *Turicibacter* abundance and AD and provide a basis for further microbiome studies of murine models for AD.

## Background

Alzheimer’s disease (AD) remains a debilitating problem for the health and wellbeing of people all over the world. Researchers are working to untangle the mystery of AD from almost every conceivable angle. We now understand much about the physiological hallmarks of the disease, including the characteristic brain changes,^1^ and many of the risk factors for AD, both genetic^2^ and environmental or lifestyle based.^3^ However, despite this extensive research, a cure remains tragically elusive. Of the six drugs currently approved for the treatment of AD, five treat the symptoms of the disease but do not slow its progression. The sole exception is the controversial monoclonal antibody-based drug “aducanumab” that is reported to reduce the accumulation of amyloid beta (Aβ) plaques.^4–7^

One reason a cure has remained elusive is an insufficient recapitulation of the neuropathology and behavioral aspects of AD in animal models. Scientists have developed more than 100 genetically engineered mouse lines that express specific aspects of AD clinicopathology, but – as valuable as they are – none of these models fully recapitulate all important aspects of the disease.^8,9^ This is especially the case for late onset AD (LOAD), which accounts for most cases.^10^ In response to this challenge, researchers have formed a National Institute on Aging (NIA) funded consortium to develop better animal models for AD.^11^

Awareness is growing around the role that host microbial community plays in AD pathology,^12–16^ including the interrelationship between inflammation, microbial exposure, and AD development.^17–19^ A slate of studies have shown that antibiotics improve behavioral symptoms and reduce amyloid plaque pathology,^14,20,21^ and germ-free murine models for AD exhibit reduced neuropathology relative to those with a normal microbiome.^22^ Human gut microbiome composition has been associated with AD in several studies^23,24^ and blood bile acid signatures (a readout of both host and microbiome metabolism) have been correlated with cognitive decline in AD.^25^ Despite this increased evidence, those developing and validating animal models often neglect to consider microbiome to be a critical component.

Considering the mounting evidence, it is imperative that we examine model animals from the perspective of the gut microbiome. Here we present the longitudinal characterization of the fecal and cecal microbiomes and metabolomes of a familial AD transgenic mouse model (5xfAD) in comparison to wild-type (WT) B6J mice. Originally introduced in 2006,^26^ the 5xfAD mouse has become the gold-standard for familial AD research and is the basis for many emerging mouse lines. 5xfAD mice exhibit many features of AD, such as intraneuronal-amyloid aggregates, neurodegeneration, neuron loss, and impaired memory, with progressive changes emerging as early as 6 weeks of age.^26^ Two recent MODEL-AD studies have comprehensively evaluated many important characteristics of the 5xfAD mouse, including behavior, cognition, and neuropathology.^27,28^ Aside from a brief assessment by Oblak *et. al.*, the microbiome was left largely undiscussed.^28^ Here we applied shotgun metagenomic sequencing and metabolomics to explore the microbes and metabolites that differ between WT and 5xfAD mice from 4-18 months of age. Associations between the microbiome and metabolome and the age, sex, and housing assignment of the mice were examined. We also examined the co-occurrences of metabolites and microbes.

## Methods

### Animal Conditions & Fecal Collection

All animal experiments were approved by the UC Irvine Institutional Animal Care and Use Committee and were conducted in compliance with all relevant ethical regulations for animal testing and research. Animals were bred and aged together in the Transgenic Mouse Facility at UCI and maintained in a 12/12-hour light/dark cycle. 5xFAD hemizygous (B6.Cg Tg(APPSwFlLon,PSEN1*M146L*L286V)6799Vas/Mmjax, Stock number 34848-JAX, MMRRC) and its wildtype littermates were produced by crossing or IVF procedures with C57BL/6 J (Jackson Laboratory, ME) females. After weaning, they were housed together with same-sex littermates until harvest. Littermate female and male mice from 4, 8, 12, and18 months of age were used in this study. All mice were fed LabDiet® Irr6f (7% fat) diet and sustained on pH 2.5 – 3.0 autoclaved water.

Immediately prior to sacrifice, the animals were isolated into individual cages for fecal collection. All working areas and tools were sterilized with 70% ethanol. Fecal pellets and cecum samples were deposited directly into 1.5 mL Eppendorf tubes. All sample tubes were placed on dry ice following collection. All samples were stored at −80°C until analysis.

### Sequence Library Preparation

DNA was extracted from both fecal and cecal samples by Zymo Research Corp. using the ZymoBIOMICS-96 MagBead DNA Kit (Zymo Research, Cat. # D4302). The quantity of DNA in each sample was measured with the Quant-iT PicoGreen dsDNA Assay Kit (ThermoFicher, Cat# P11496) read with a Synergy H1 Microplate reader (BioTek, Cat # BTH1M). Sequence libraries were prepared from the extracted DNA samples using the Nextera DNA Flex Library Prep Kit (Illumina, Cat. # 20018705) following a low volume variation of the standard protocol.^29^

The average input DNA for each sample was 391.6 ng (std dev = 54.8 ng). Samples were prepared for PCR with Kapa HiFi HotStart ReadyMix (Roche, Cat # 07958935001) using the following primers:

KAPA-PCR-F: 5’ – AATGATACGGCGACCACCG*A – 3’

KAPA-PCR-R: 5’ – CAAGCAGAAGACGGCATACG*A – 3’

PCR was performed with an Eppendorf Mastercycler Nexus Gradient (Eppendorf, Cat # 2231000665) using the standard thermal cycles for the Nextera Flex kit. The resulting sequence fragments were analyzed on an Agilent Bioanalyzer to determine fragment length distribution (Agilent, Cat # G2939BA). Sequence libraries were pooled based on DNA concentration as determined by the Bioanalyzer. After pooling, the libraries were sent to Novogene Co., Ltd. for sequencing on an Illumina HiSeq4000.

### Sequence data processing

Sequence libraries were downloaded from the Novogene FTP website. The libraries consisted of 456 samples pooled into a single dual-index paired-end library and run across two lanes. The two paired-end aggregate FASTQ files were parsed using BBDuk to trim adapters and remove artifacts, mouse DNA (GCA_000001635.8), and rat DNA (GCA_000001895.4). The cleaned files were demultiplexed using demuxbyname.sh inside of the BBmap suite. To assign microbial taxonomy, GenBank’s NT database (version 1/29/20) was retrieved using the update_blastdb PERL script. Each pair of metagenome files was queried against NT using GenBank’s MEGABLAST algorithm from the BLAST+ package (version 2.10.0) using all default parameters. For phylogenic assignment, the NCBI Taxonomy database dump files were downloaded from the NCBI FTP site. BLAST outputs were parsed to retrieve the accession numbers of the hits, and then GenBank’s blastdbcmd program, along with the taxonomy dump files, were used to get the full taxonomic rank of each hit.

When a sequence read maps to multiple taxon, it is acceptable to either ignore all hits for that read or divide the hit evenly across the taxa. The first option excludes taxa with homologous sequences, whereas the second option introduces false positives into the taxonomical assignments and alters abundance estimates. We choose the second option and mitigated false positives by excluding taxa with abundance counts less than one.

During analysis, a major update was made to *Lactobacillus* taxonomical assignments which included the parsing of *Lactobacillus* into 25 new genera.^30^ To accommodate this change, we updated the names of our final species-level taxonomical assignments using the NCBI Taxonomy Browser (https://www.ncbi.nlm.nih.gov/Taxonomy/TaxIdentifier/tax_identifier.cgi).

### Plasma Metabolomics

Approximately 1.5 mL of blood was drawn after cessation of respiration via a closed-chest cardiac blood draw, using a tuberculin syringe (1.0 mL) and a 25-gauge needle. Aspirated blood was immediately transferred to a 2.0 mL EDTA tube, at a rate of approximately 1 mL/30 seconds, to mitigate hemolysis. The EDTA tube was inverted 10 times and placed in wet ice. EDTA blood tubes were processed within 1hour of placement into ice. Blood components were separated via centrifugation at 2600 RPM (1500 x g) for 10 minutes at 20°C. Plasma was carefully aspirated from the EDTA tube using a micropipette, and aliquoted as 100 microliter (μL) volumes into siliconized cryovials and stored at −80°C until metabolomic analysis. The small plasma volumes allowed for a single freeze-thaw cycle prior to metabolomic analysis.

We performed targeted metabolomic measurements using Stable Isotope Dilution, Multiple Reaction Monitoring, Mass Spectrometry (SID-MRM-MS). We used the Biocrates AbsoluteIDQ® P180 (BIOCRATES, Life Science AG, Innsbruck, Austria) kit as in our previous work.^31–33^ For this targeted analysis, frozen plasma samples were thawed on ice, vortexed, and processed as per the manufacturer’s instructions. The metabolite abundances were measured in a 96-well format, including seven calibration standards and three quality control samples integrated as part of the kit. The kit was run on a triple quadrupole mass spectrometer (Xevo TQ-S, Waters Corporation, USA) operating in the multiple reaction monitoring (MRM) mode. The kit provided absolute quantitation of 21 amino acids, hexoses, carnitines, 39 acylcarnitines, 15 sphingomyelins, 90 phosphatidylcholines, and 19 biogenic amines. The flow injection analysis (FIA) tandem mass spectrometry (MS/MS) method is used to quantify the 144 lipids simultaneously by multiple reaction monitoring. The other metabolites are resolved on the Ultra Performance Liquid Chromatography (UPLC) and quantified using scheduled MRMs. Each metabolite’s abundance was calculated from area under the curve by normalizing to the respective isotope labeled internal standard provided with the kit. The concentration of each metabolite is expressed in nmol/L. We used EDTA plasma samples spiked with standard metabolites as quality control samples to assess reproducibility of the assay. The quantitated data were uploaded to RStudio and values < LOD were set to NA or 0, as appropriate for the specific analysis.

### Fecal Metabolomics

Fecal material was shipped on dry ice to the West Coast Metabolomics Center (WCMC) at UC Davis for extraction and analysis via hydrophilic interaction chromatography (HILIC) time-of-flight (TOF) mass spectrometry (MS). Standardized WCMC extraction and analysis protocols for “Biogenic amines by HILIC-QTOF MS/MS” were used throughout (analogous to those described in Ding 2021). Approximately 4 mg of fecal material was used from each sample, the final extract was dissolved in 100 μL acetonitrile/water (4:1, v/v) containing internal control standards, and 3 μL of each extract was analyzed on a SCIEX 6600 TTOF using both positive and negative ionization. We received the resulting data as a table containing ion identifications (where possible) and peak heights normalized to the sum of the internal standards.

### Microbiome analysis

Processed sequence data were analyzed using RStudio Version 1.4.1106^34^ without rarefaction. Most graphics were created using the R package ggplot2.^35^ The heatmaps in **Figure 1** were generated from standard normalized abundance data using the pheatmap package with “average” clustering.^36^ Shannon diversity, nonmetric multidimensional scaling (NMDS), and permutational multivariate analysis of variance (PERMANOVA) were computed using the Vegan package.^37^ Both NMDS and PERMANOVA were applied to the Bray-Curtis dissimilarity matrix. PERMANOVA was performed using the Adonis function with 999 permutation and the following formula: formula = data ~ Genotype + Sex + SampleType + Cohort + HousingID. Principal coordinate analysis (PCoA) was performed on the Bray-Curtis dissimilarities of the relative abundances using the cmdscale function from the stats package, and the resulting data were visualized with ggplot2.^35^ Random forest was performed using the rfPermute package (v2.1.81). Linear mixed-effects model (LME) analysis was performed on relative abundance data using the lmer function in the lme4 package^38^ with an abundance threshold of 0.001 and Housing ID as the random effect.

**Figure 1.**
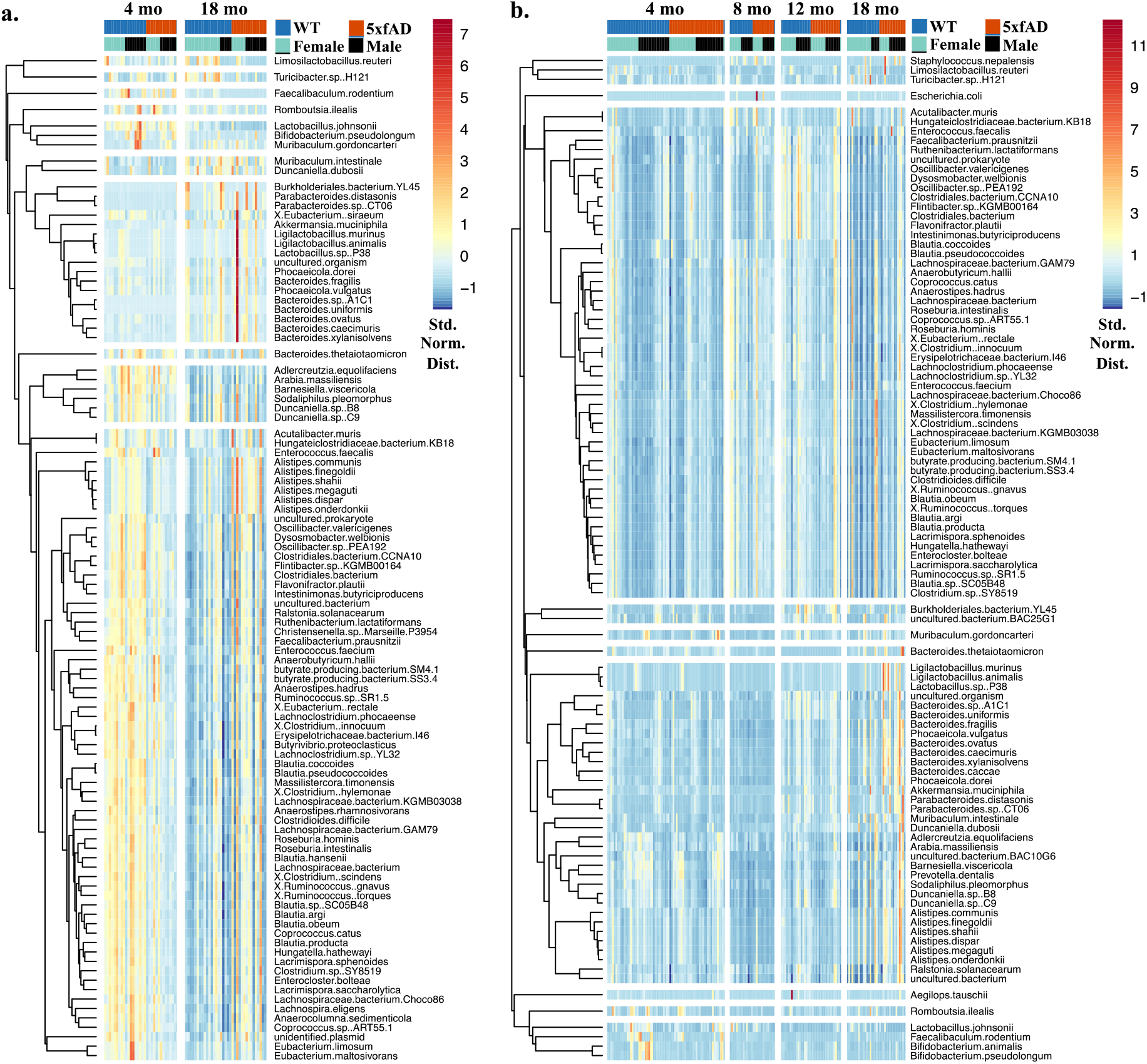
Clustered heatmaps of the 100 most abundant microbial species in the cecal (**a**) and fecal (**b**) samples with each sample represented by a column and each species as a row. The colour scale indicates abundance (standard normal distribution), with hierarchical clustering used to group microbes with similar abundance trends.

### Turicibacter *analysis with Bowtie 2*

To determine the abundance of *Turicibacter* in our samples, three reference genomes were downloaded from NCBI: *Turicibacter sp.* H121 (GenBank assembly accession: GCA_001543345.1), *Turicibacter sp.* HGF1 (GenBank assembly accession: GCA_000191865.2), and *Turicibacter sanguinis* (GenBank assembly accession: GCA_013046825.1). Bowtie2 v2.4.4 was used with the default parameters to align reads from our samples to each *Turicibacter* reference genome. *Turicibacter* reads per sample were obtained and converted to relative abundances using the total number of reads per sample. A Wilcoxon test was used for significance testing in R v3.6.

### Metabolomics data analysis

Analysis of both the fecal and plasma metabolomics data was performed using methods adapted from the microbiome analysis described above. Spearman’s correlations between metabolites and species were generated using the cor function (method = spearman) from the stats package, and the resulting data were plotted using the pheatmap function from the pheatmap package.^36^

## Results

### Shotgun Metagenomic Sequencing Sample Demographics

Shotgun metagenomics was performed on 66 cecal samples from 4- and 18-month-old mice and 170 fecal samples from 4-, 8-, 12-, and 18-month-old mice. The samples had a slightly higher representation from WT than 5xfAD (128 WT vs 108 5xfAD) and slightly more females than males (124 females vs 114 males). The average number of reads per sample for each cohort ranged from 1.1 – 1.9 million, with 18 – 28% of those reads aligning to a genome in GenBank’s NT database. See **Table S1** for a more thorough demographic breakdown.

### Overview of Most Abundant Microbes

Clustered heatmaps of the 100 most abundant microbial species in cecal (**Figure 1a**) and fecal (**Figure 1b**) samples showed that many species differed in abundance with respect to both age and genotype. For example, the first three bacteria in the cecal sample heatmap (*Limosilactobacillus reuteri*, *Turicibacter* sp. H121, and *Faecalibaculum rodentium*) had lower abundances in the 18 month 5xfAD samples relative to both the 18 month WT and 4 month samples. Those microbes in the 7^th^ cecal cluster (comprised of 17 species from the genera *Burkholderiales*, *Akkermansia*, *Ligilactobacillus*, and *Bacteroides*, among others) were generally more abundant at 18 months than 4 months. In the fecal samples (**Figure 1b**), *T.* sp. H121 and *L. reuteri* were found together in the first cluster with *S. nepalensis*, while *F. rodentium* is found in the third cluster. Organisms in the 7^th^ cluster of the fecal samples had increased abundances at 18 months of age. Among others, this cluster included multiple species from the genera *Ligilactobacillus*, *Bacteroides*, *Alistipes*, and *Duncaniella*. Differences with respect to sex were not readily apparent in this representation of the data.

### Alpha and Beta Diversity

Examination of the richness (total number of microbial species; **Figure 2a**) and Shannon diversity (a measure of both richness and relative abundance; **Figure 2b**) of the fecal microbiomes showed significant overall differences with respect to age but not genotype (by two-way ANOVA). Both the richness and Shannon diversity of the fecal microbiome increased between 4 and 8 months of age, and then successively decreased. The richness and the Shannon diversity of the cecal microbiomes did not significantly differ with respect to age or genotype (p > 0.05 by Kruskal-Wallis 1-way ANOVA). Separating the cecal and fecal data based on sex (*i.e.,* evaluating males and females separately; **Figure S1**) also failed to uncover significant differences in Shannon diversity by genotype.

**Figure 2.**
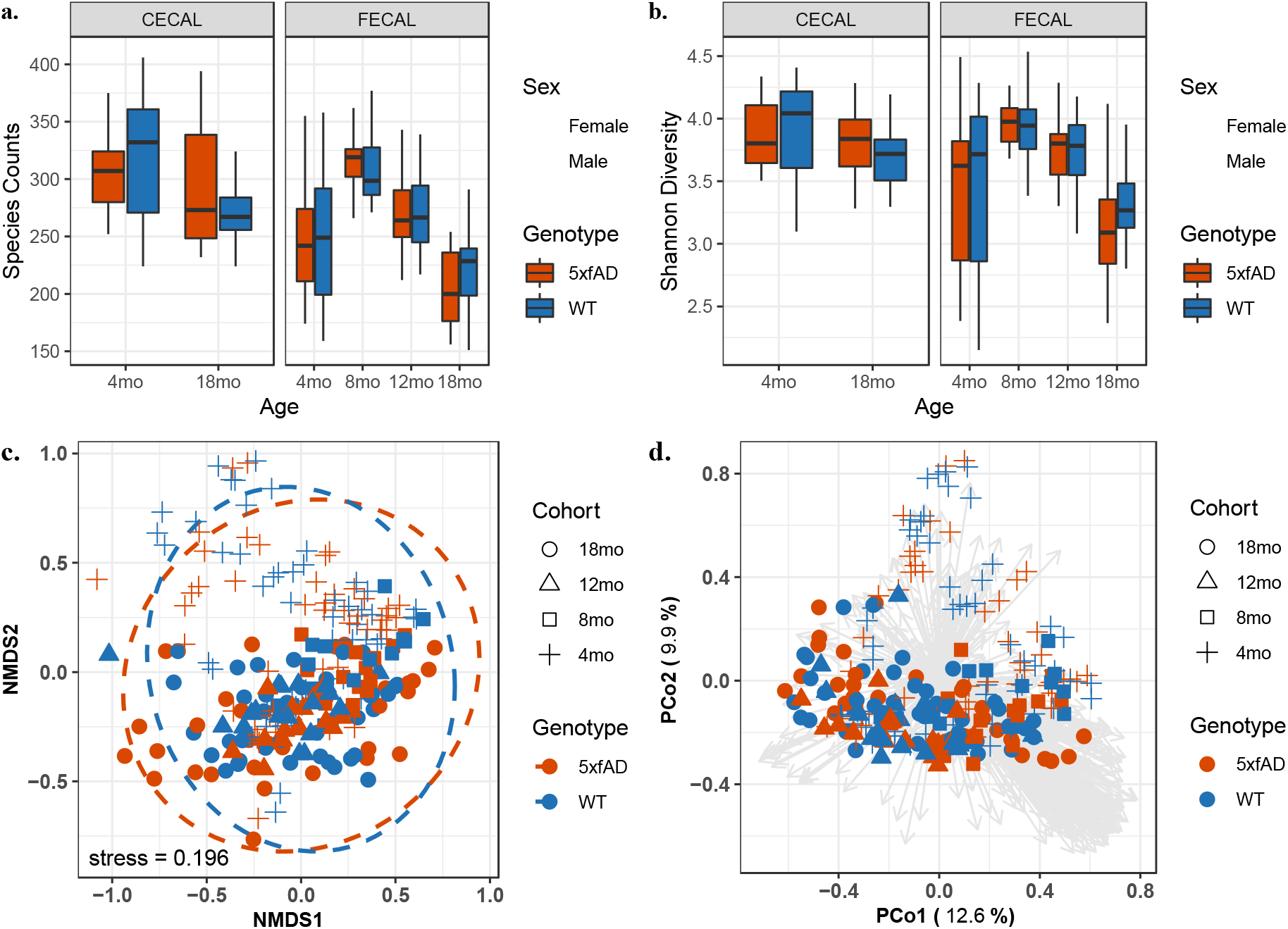
Alpha and beta diversities of the cecal and fecal microbiomes of 5xfAD and WT mice from 4 to 18 months of age. Species richness (**a**) and Shannon diversity (**b**) did not significantly differ between genotypes but did significantly differ between age groups (richness: p < 0.05 for age, p > 0.05 for genotype by 2-way ANOVA; Shannon p < 0.05 for age, p > 0.05 for genotype by Kruskal-Wallis 1-way ANOVA). Beta diversity as visualized by non-metric multidimensional scaling (NMDS) (**c**) and principal coordinate analysis (PCoA). With both methods separation was generally observable by age but not genotype. The box plots in (**a-b**) show the median value (computed from absolute abundance) +/-the first and third quartiles, with the whiskers showing the range or 1.5 x the interquartile range, whichever is less. The analysis in (**c-d**) was performed on relative abundance data, with ellipses in **c** representing the 95% confidence interval for each genotype.

Beta diversity – as evaluated through both non-metric multidimensional scaling (NMDS; **Figure 2c**) and principal coordinates analysis (PCoA; **Figure 2d**) of the Bray-Curtis dissimilarity matrix – showed general separation by age but not genotype. The NMDS results exhibited a high stress value (0.196), and nearly complete overlap between the 95% confidence intervals for genotype, indicating weak association. Inspection of the PCoA loadings (grey vectors in **Figure 2d**) revealed separation arising from contributions of many species (as opposed to a few species driving differences between groups). PCoA of the cecal and fecal data separately (**Figure S2a-b**) and males and females separately (**Figure S2b-c**) showed more distinct separation by age, but not by genotype.

Permutational multivariate analysis of variance (PERMANOVA) was performed on cecal and fecal samples together, cecal samples only, and fecal samples only (**Table 1**). Significant differences (p < 0.05) in microbiome composition were observed between genotypes, sexes, sample types, age groups, and housing IDs, with the largest proportion of the variance (R^2^ > 57% in all cases) explained by housing ID. Age also accounted for a substantial portion of the variance (R^2^ > 16% in all cases). Sex and genotype accounted for more of the variance in the cecal samples than the fecal samples. An important point to note is that many of the variables used in this comparison are inextricably conflated as: (1) littermates of the same sex were often (but not always) housed together; (2) males and females were always housed separately; and (3) mice of different age groups were housed separately.

**Table 1.** Permutational multivariate analysis of variance (PERMANOVA) of the model animal microbiomes. Shown are separate calculations for all samples (cecal and fecal combined), cecal samples only, and fecal samples only. Bacterial species composition was significantly affected by all variables examined, including genotype. Many variables were confounding, as littermates of the same sex often (but not always) shared the same cage.

All Samples
|  | Df | SumsOfSqs | MeanSqs | F.Model | R2 | Pr(>F) |
| --- | --- | --- | --- | --- | --- | --- |
| Genotype | 1 | 0.1614 | 0.16139 | 5.667 | 0.00596 | 0.001 |
| Sex | 1 | 0.7444 | 0.74441 | 26.140 | 0.02748 | 0.001 |
| Sample Type | 1 | 1.2861 | 1.28607 | 0.04748 | 0.04748 | 0.001 |
| Age | 3 | 4.3819 | 1.46063 | 0.16177 | 0.16177 | 0.001 |
| Housing ID | 51 | 15.4449 | 0.30284 | 10.634 | 0.57018 | 0.001 |
| Residuals | 178 | 5.0690 | 0.02848 |  | 0.18713 |  |
| Total | 235 | 27.0877 |  |  | 1.00000 |  |

Cecal Only
|  | Df | SumsOfSqs | MeanSqs | F.Model | R2 | Pr(>F) |
| --- | --- | --- | --- | --- | --- | --- |
| Genotype | 1 | 0.1314 | 0.13142 | 6.027 | 0.02311 | 0.001 |
| Sex | 1 | 0.2619 | 0.26192 | 12.012 | 0.04605 | 0.001 |
| Age | 1 | 0.9925 | 0.99254 | 45.517 | 0.17450 | 0.001 |
| Housing ID | 21 | 3.4078 | 0.16228 | 7.442 | 0.59915 | 0.001 |
| Residuals | 41 | 0.8940 | 0.02181 |  | 0.15719 |  |
| Total | 65 | 5.6878 |  |  | 1.00000 |  |

Fecal Only
|  | Df | SumsOfSqs | MeanSqs | F.Model | R2 | Pr(>F) |
| --- | --- | --- | --- | --- | --- | --- |
| Genotype | 1 | 0.0782 | 0.07823 | 2.489 | 0.00389 | 0.023 |
| Sex | 1 | 0.5058 | 0.50583 | 16.090 | 0.02515 | 0.001 |
| Age | 3 | 3.3976 | 1.13253 | 36.025 | 0.16892 | 0.001 |
| Housing ID | 51 | 12.5791 | 0.24665 | 7.846 | 0.62541 | 0.001 |
| Residuals | 113 | 3.5524 | 0.03144 |  | 0.17662 |  |
| Total | 169 | 20.1132 |  |  | 1.00000 |  |

### Microbial species whose abundances differed between 5xfAD and WT animals

Random forest was used to evaluate differences in the microbiome between genotypes (**Figure 3**). AD pathology becomes more pronounced as the mice age, therefore 18 month mice were the primary focus of the random forest analysis. Clear separation was observed between genotypes both when the cecal and fecal sample types were combined (**Figure 3a**) and when they were considered separately (**Figure 3b**). In both cases, overlapping ellipses (which represent the 95% confidence intervals) showed that the groups share many species in common. *Turicibacter* sp. H121 was the species most responsible for the random forest separation (> 10% mean decreased accuracy in both separations), with *Romboutsia ilealis* and *Lactobacillus johnsonii* also playing substantial roles (**Figure 3c–d**).

**Figure 3:**
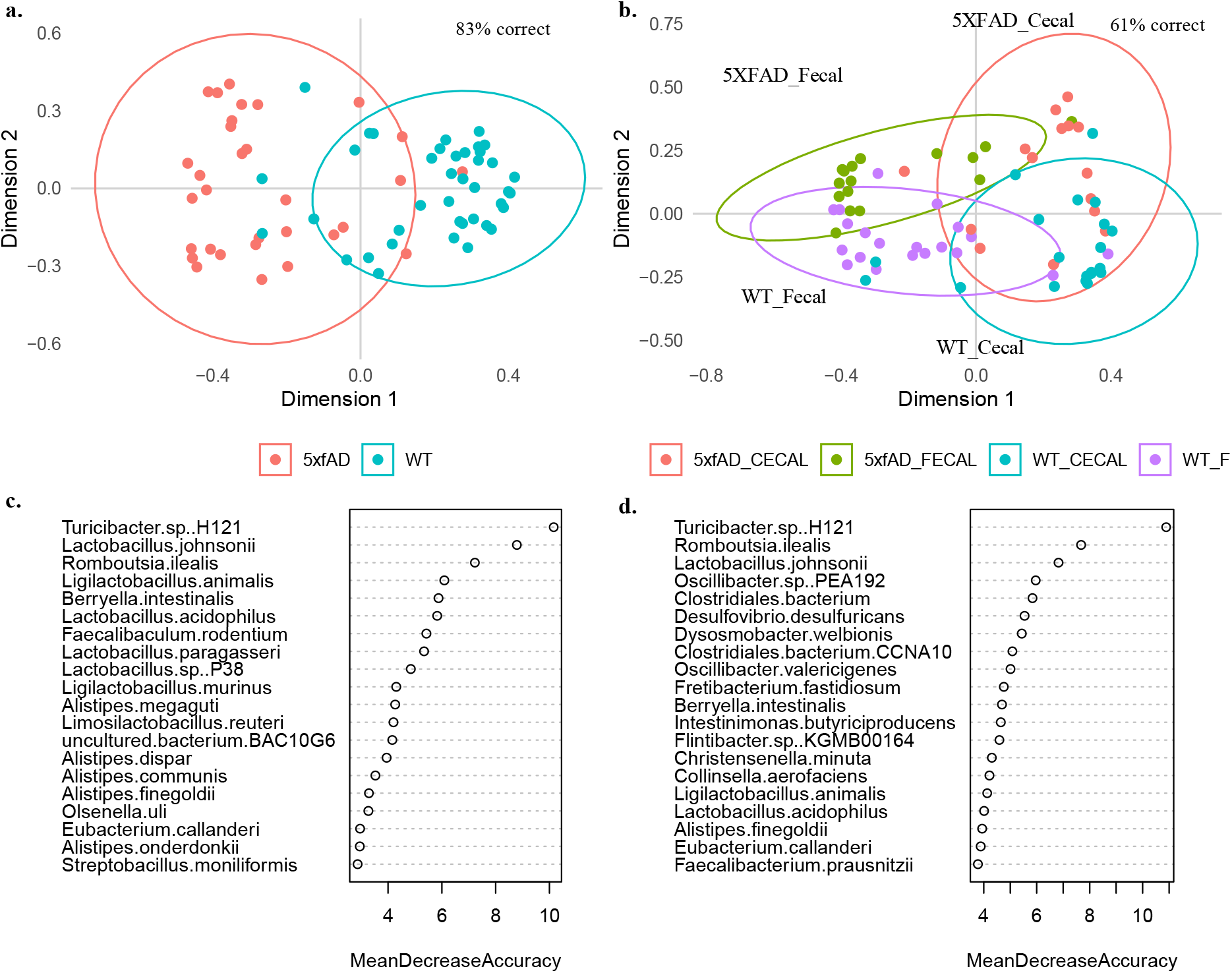
Random forest analysis of cecal and fecal microbiomes from 18 month animals. Separation by genotype only (**a**, **c**) and separation by genotype and sample type (**b**, **d**). *Turicibacter* sp. H121 was responsible for the largest proportion of separation in both analyses.

Random forest of the 18 month samples with grouping by sample type and sex (ignoring genotype differences) also showed interesting trends (**Figure S3a-d**). Grouping by sample type showed an out-of-bag accuracy of 90% with *Oscillibacter* sp. PEA192, *Flavonifractor plautii*, and *Desulfovibrio desulfuricans* contributing heavily to the separation (**Figure S3a-b**). Grouping by sex was 97% accurate, with *Limosilactobacillus reuteri* and *Lactobacillus johnsonii* playing the largest role in the separation (**Figure S3c-d**). Differences with respect to age were also examined (**Figure S3e-f**), with large overlap between each age cohort. The 4 month samples showed the most variation and the least overlap with the other age cohorts. *Staphylococcus nepalensis* was the species most responsible for separation by age, with a mean decreased accuracy (MDA) of almost 12%.

Random forest is normally applied as a classification method and can lead to false positives when used to identify differentially abundant species. Therefore, we also applied a linear mixed effects (LME) model to identify bacterial species whose abundances significantly differed between the WT and 5xfAD genotypes. The model was performed on all samples in aggregate [*i.e.*, 5xfAD (all age groups & sample types) vs WT (all age groups & sample types)] as well as to each age group and sample type individually. For example, the comparisons made for the 18 month samples were: (1) 5xfAD fecal and cecal vs WT cecal and fecal; (2) 5xfAD fecal vs WT fecal; and (3) 5xfAD cecal vs WT cecal. After correcting for the false discovery rate, significantly different species were only found in the 18 month samples. The 18 month cecal samples had 15 differentially abundant species (padj < 0.05) whereas the other two groupings each had six differentially abundant species (**Table 2**). Five species were significant across all three groupings: *Turicibacter* sp. H121, *Romboutsia ilealis*, *Lactobacillus* sp. P38, *Ligilactobacillus murinus*, and *Ligilactobacillus animalis*. Among others, six species from the genus *Alistipes* were differentially abundant in the 18 month cecal samples only. A large proportion of the species (11/17) were also among the top 20 species contributing to random forest separation shown in Figure 3.

**Table 2:** Species whose abundances significantly differed between 18 month WT and 5xfAD mice with Housing ID as the random variable. The “5xfAD Abn.” column indicates whether the species was found in relatively higher or lower abundance in the 5xfAD samples compared to the WT. Species in purple were found to be significantly different in all three comparisons and species with an asterisk were among the top drivers of separation in the random forest analysis shown in **Figure 2a** and **2c**. LME computed p-values (padj) are false-discovery rate (FDR) corrected.

| 18 Month Fecal & Cecal Samples |  |  | 18 Month Cecal Samples |  |  |
| --- | --- | --- | --- | --- | --- |
| Species | $p_{adj}$ | 5xfAD Abn. | Species | $p_{adj}$ | 5xfAD Abn. |
| *Turicibacter* sp. H121* | <5E-7 | Lower | *Turicibacter* sp. H121* | <5E-3 | Lower |
| *Romboutsia ilealis* * | <5E-4 | Lower | *Romboutsia ilealis* * | <0.05 | Lower |
| *Lactobacillus* sp. P38* | <5E-4 | Higher | *Alistipes megaguti* * | <0.05 | Higher |
| *Ligilactobacillus murinus* * | <5E-4 | Higher | *Alistipes dispar* * | <0.05 | Higher |
| *Ligilactobacillus animalis* * | <5E-4 | Higher | *Alistipes onderdonkii* * | <0.05 | Higher |
| *Burkholderiales* bacterium YL45 | <5E-3 | Lower | *Alistipes shahii* | <0.05 | Higher |
| 18 Month Fecal Samples |  |  | *Ligilactobacillus murinus* * | <0.05 | Higher |
| Species | $p_{adj}$ | 5xfAD Abn. | *Lactobacillus* sp. P38* | <0.05 | Higher |
| *Turicibacter* sp. H121* | <5E-3 | Lower | *Alistipes finegoldii* * | <0.05 | Higher |
| *Ligilactobacillus murinus* * | <0.05 | Higher | *Duncaniella* sp. C9 | <0.05 | Lower |
| *Lactobacillus* sp. P38* | <0.05 | Higher | *Ligilactobacillus animalis* * | <0.05 | Higher |
| *Ligilactobacillus animalis* * | <0.05 | Higher | *Alistipes communis* * | <0.05 | Higher |
| *Romboutsia ilealis* * | <0.05 | Lower | *Duncaniella* sp. B8 | <0.05 | Lower |
| *Adlercreutzia equolifaciens* | <0.05 | Lower | *Muribaculum intestinale* | <0.05 | Lower |
|  |  |  | *Lactobacillus johnsonii* * | <0.05 | Lower |

Relative abundance plots for all 17 differentially abundant species can be found in **Figure S4**, with faceting by sample type and sex to allow inspection of more subtle abundance differences. Abundances of 8 of the 17 significant species were lower in 18 month 5xfAD mice, and 9 of the 17 species were present at a higher abundance. With the exception of *Ligilactobacillus murinus* (which had relative abundances as high as 50%) the differentially abundant species were relatively minor contributors to the overall microbiome composition, with relative abundances below 2.5%.

### *Multiple* Turicibacter *species were significantly less abundant in 18 month 5xfAD animals*

*Turicibacter* was previously identified as a potential mediator of the gut-brain axis in AD through its ability to consume, and regulate the production of, 5-hydroxytryptamine (serotonin).^39^ Further studies have highlighted the potential role of serotonin in AD (recently reviewed by Aaldijk *et. al.* in 2022).^40^ These previous findings, as well as the importance of this genus in our random forest and LME analysis, motivated us to persue a more detailed investigation. A Bowtie 2 sequence alignment was performed against reference genomes for three species of *Turicibacter, T.* sp. H121, *T.* sp. HGF1, and *T. sanguinis*, that encode a neurotransmitter sodium symporter-related protein with sequence and structural homology to mammalian serotonin transporter (SERT; see **Methods** for more details). All three species were present in both the fecal and cecal microbiomes (**Figure 4a–b**). In agreement with the NCBI Blast data for *T.* sp. H121 shown in **Figure S4**, all three species were significantly less abundant in 5xfAD mice relative to the WT mice at 18 months of age (p < 0.05). Any apparent differences at other ages were not statistically significant.

**Figure 4.**
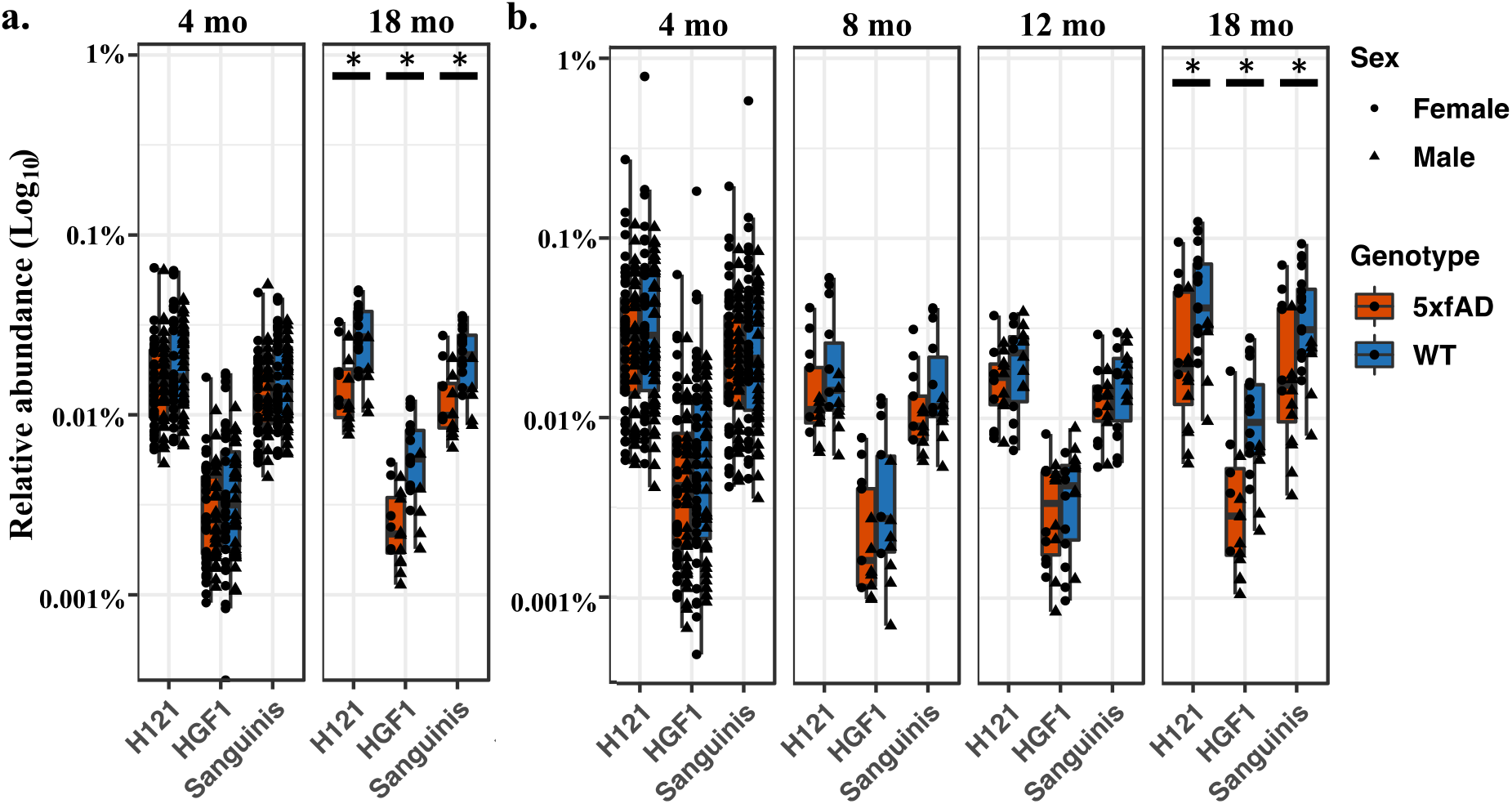
Relative abundances for *Turicibacter* spp. in fecal (**a**) and cecal (**b**) samples generated via Bowtie2. Shown are *T. sanguinis*, *T*. sp. H121, and *T*. sp. HGF1. The WT and 5xfAD cohorts in (**a-b**) are significantly different (p < 0.05 via the Wilcox rank sum test) for all three *Turicibacter* species at 18 months of age only.

### Plasma Metabolomics Reveals Differentially Abundant Phospholipids and Amino Acids with Respect to Age, Sex, and Genotype

Quantitative metabolomics was performed on the blood plasma of the 12 and 18 month mice in our study. For this data set we utilized the Biocrates Absolute-IDQ P180 Kit (see **Methods**), which was previously applied to develop blood-based biomarker assays for preclinical AD.^31–33^ From this list of 188 potential analytes, we received concentrations for 137 metabolites within the assay’s quantitation limits for more than 50% of our samples, which was the quality threshold used for this report.

The metabolites showed similar aggregate concentrations for 12 and 18 months of age (p > 0.05 by Kruskal-Wallis test) (**Figure S5a**). Shannon alpha diversity was also similar across all age and genotype groupings (p > 0.05 by Kruskal-Wallis test) (**Figure S5b**). PCoA did not reveal clear separation with respect to age, sex, or genotype (**Figure S5c**). PERMANOVA of the Bray-Curtis dissimilarity matrix (**Figure S5d**) showed sex to be the only significant distinguishing factor (p = 0.037; R^2^ = 0.069). Housing ID accounted for almost 40% of the variance in the data set but was not significant (p = 0.357; R^2^ = 0.39).

A clustered heatmap of all 137 validated plasma metabolites **(Figure 5a**) shows metabolite specific differences based on age, genotype, and sex. Relative to their 12 month counterparts, the 18 month mice appeared to have lower concentrations of α-aminoadipic acid, serotonin, glutamic acid, aspartic acid, spermidine, and spermine. Most detected phosphatidylcholines (PCs) [both diacyl (aa) PCs and acyl-alkyl (ae] PCs) were depleted in 18 month 5xfAD mice, while many amino acids (shown primarily in the 6^th^ cluster) had an increased relative concentration. Sex-specific differences were pronounced for most detected PCs, which clustered together in the 4^th^ cluster.

**Figure 5.**
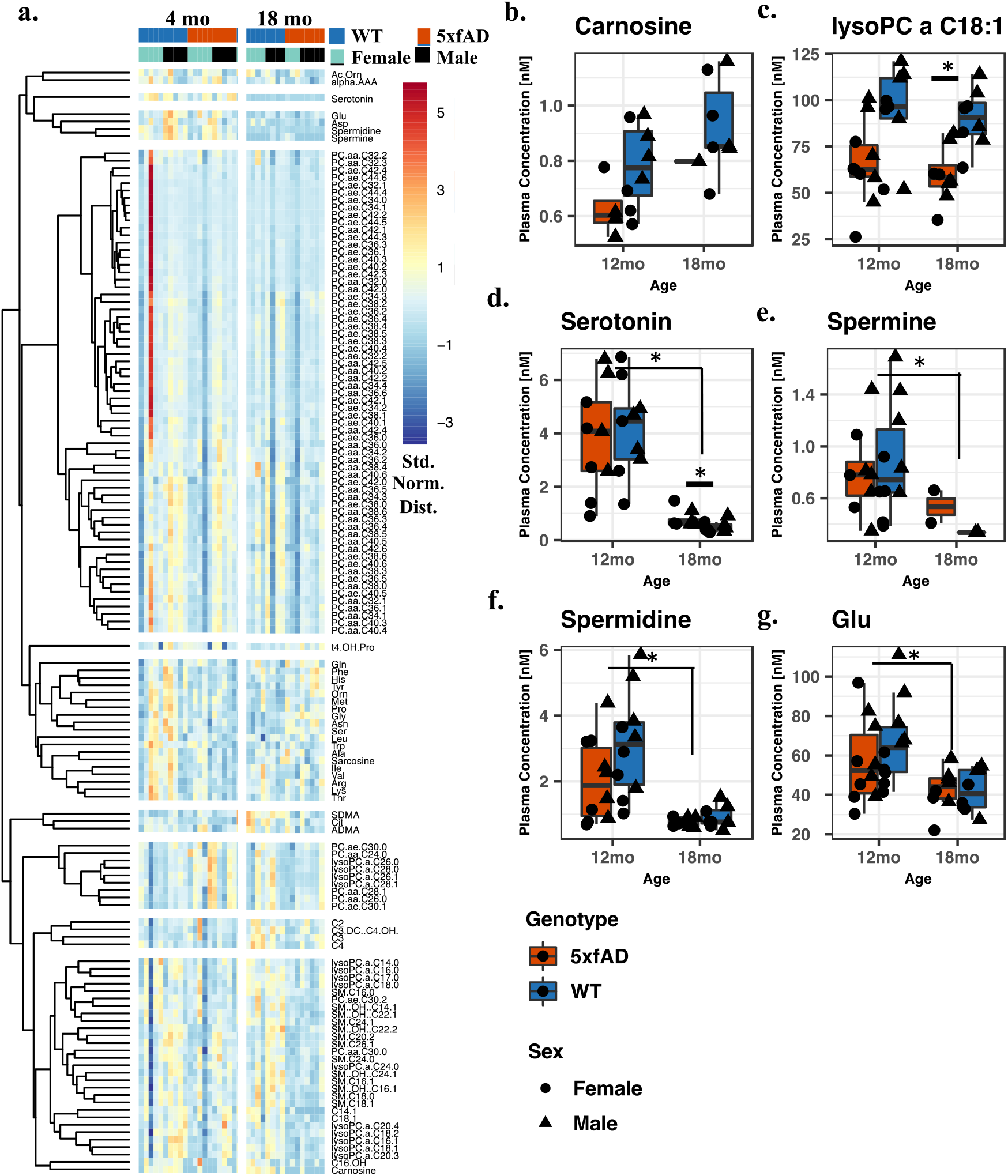
Quantitative plasma metabolomics. (**a**) Heatmap of the 137 quantified metabolites in the plasma of 12 and 18 month old mice. Shown is the standard normal abundance distribution, with hierarchical clustering to group metabolites with similar abundance trends. (**b-c**) Metabolites with significantly different concentrations with respect to genotype (padj < 0.05 by LME). (**d-g**) Metabolites with significantly different concentrations with respect to age (padj < 0.05 by LME). The asterisks in **b-g** indicate metabolites found significantly differ in each comparison by the Wilcoxon signed rank test (p<0.05).

Random forest (**Figure S6**) showed separation by age (75% out-of-bag accuracy), sex (78% accurate), and genotype (81% accurate for 18 month samples only). Serotonin was the metabolite most responsible for age related separation (MDA = 8.5%), closely followed by spermidine (MDA = 7%) and spermine (MDA = 7%). Separation with respect to sex was dominated by sphingomyelin sphingolipids (SMs) and phosphatidylcholines (PCs) from both the diacyl (aa) and acyl-alkyl (ae) subclasses. SM C20:2, PC aa C40:3, PC aa C42:6, PC ae C42:0, PC aa C34:3, PC ae C34:2, SM (OH) C22:2 topped this list, each accounting for more than 5% MDA. Random forest separation with respect to genotype was negligible when both age groups were considered together (61% accurate). The most predictive molecules for genotype at 18 months of age were glycine (MDA = 7%), carnosine (MDA = 4.5%), serine (MDA = 4.5), SM C24:1 (MDA = 4.2%) and serotonin (MDA = 4%).

LME was also performed using housing ID as the random effect and several significantly different metabolites (padj < 0.05) were observed with respect to age, sex, and genotype (see **Table S2**). The concentrations of the metabolites that significantly differed between genotypes are shown in **Figures 5b–c**, between ages in **Figures 5d–g**, and between sexes in **Figure S7**. Carnosine and lysoPC a C18:1 had significantly lower plasma concentrations in 5xfAD mice relative to WT (padj < 0.05). Serotonin, spermine, spermidine, and PC aa C36:0, each had significantly lower concentrations in 18 month mice relative to the younger cohort. Concentrations of two SM lipids, two PC ae lipids, six PC aa lipids, and lysine significantly differed between sexes. Separately, a targeted analysis using the Wilcoxon signed rank test revealed serotonin to be more concentrated in 18 month 5xfAD animals relative to their age-matched counterparts (**Figure S8**; p = 0.038).

### Correlations between the microbiome and the plasma metabolome

Spearman correlations were generated to better evaluate the relationships between the significant microbes at 18 months of age and the plasma metabolome. Spearman correlations between all 137 plasma metabolites and the 17 significant microbes for 18 month mice are provided **Figure 6**. **Figures S9** and **S10** show analogous correlations for all 137 plasma metabolites and the 100 most abundant microbes.

**Figure 6.**
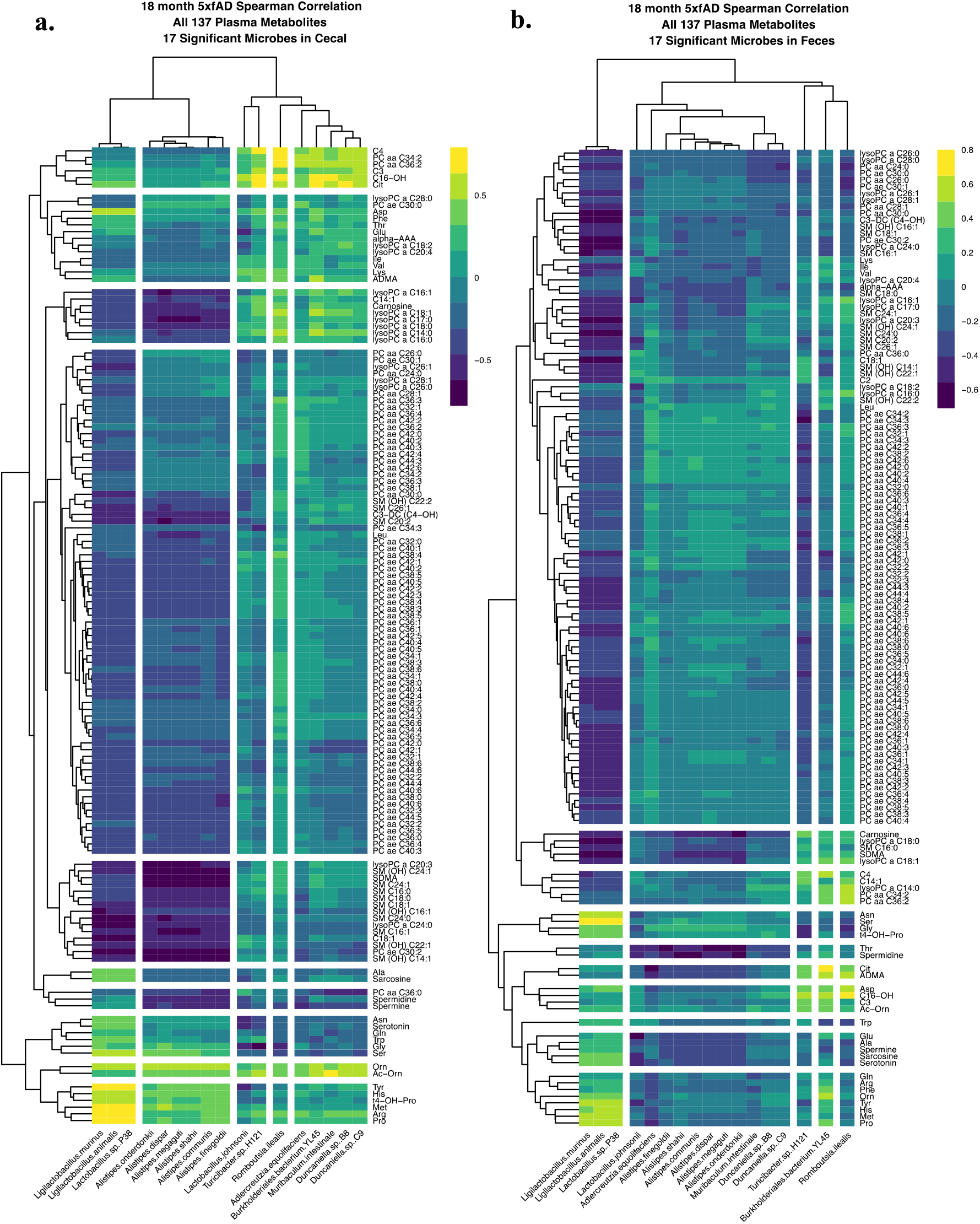
Spearman correlations between the 17 significant microbial species and all 137 quantified plasma metabolites at 18 months of age: (**a**) cecal; and (**b**) fecal samples.

For the cecal samples (**Figure 6a**), strong correlations were observed between the first cluster of bacteria (*Ligilactobacillus murinus*, *Ligilactobacillus animalis*, and *Lactobacillus* sp. P38) and PC ae C30:0, ornithine, acetylornithine, tyrosine, histidine, methionine, arginine, and proline (among others). These bacteria were anticorrelated with most lipids. Similarly, the six *Alistipes* species (which were found together in the second microbial cluster) weakly co-occurred with several amino acids in the last two metabolite groupings. The remaining 8 microbes (which were all at reduced relative abundance in the 18 month 5xfAD samples) showed correlation trends similar to one another. They were weakly anticorrelated with most metabolites aside from three acylcarnitines (propionylcarnitine (C3), butyrylcarnitine (C4), exadecenoylcarnitine (C16-OH)), PC aa C34:2, PC aa C36:2, citrulline, and acetylornithine.

All l7 microbes in the fecal correlation (**Figure 6b**), were either anticorrelated or weakly correlated with the top group of 102 metabolites. *Ligilactobacillus murinus*, *Ligilactobacillus animalis*, and *Lactobacillus* sp. P38 were correlated with the most amino acids and many acylcarnitines and strongly anticorrelated with most lipids. The six *Alistipes* species, *Muribaculum intestinale*, *Duncaniella* sp. B8 and *Duncaniella* sp. C9 were weakly correlated with or anticorrelated with all 137 metabolites. *Turicibacter* sp. H121 and *Romboutsia ilealis* were weakly correlated with many amino acids, acylcarnitines, and biogenic amines, and differed from *Burkholderiales bacterium* YL45 based primarily on the last cluster of 8 amino acids.

Untargeted metabolomics (via HILIC-QTOF-MS) was also performed on a subset of fecal samples with adequate material, namely n = 40 from the 4, 8, and 12 month old mice. Unfortunately, there was insufficient material to perform both metabolomics and genomics for the 18 month mice, and therefore there are no metabolomics data for this age cohort. Overall trends did not reveal major differences with respect to genotype. Please see the **Supplemental Results** for a more thorough review of this dataset.

## Discussion

### Overview of Microbiome Composition and Changes

Overall, the richness and Shannon diversity of the fecal microbiomes increased between 4 and 8 months and declined thereafter (**Figure 2**). Among other factors, changes in gut microbial diversity have been associated with age,^41^ sex,^41^ obesity,^42^ and general health.^43^ As revealed by PERMANOVA (**Table 2)**, most of the variance in the fecal and cecal microbiomes was attributable to the specific cage where the animals resided, and relatively minor shifts in the microbiome composition were observed when comparing factors such as the age and sex of the mice. The strong influence of cage on the microbiome is consistent with results from other studies of fecal microbiomes in 5xfAD mice.^19,28^ Such “cage effects” are routinely observed in microbiome studies, increasing variability, and complicating the association between trends in the microbial population and the experimental factors of interest.^44–47^ The consistency of these findings emphasizes the necessity of thoroughly documenting housing protocols and accounting for cage effects in experimental design (e.g., by housing animals from the control and experimental cohorts together as we did here).

### Differentially Abundant Species

Comparing the microbiome composition of 5xfAD mice to WT B6J controls revealed differences later in life (**Figures 1 and 3)**. The abundances of 17 species of bacteria were found significantly differ between genotypes at 18 months (15 species in the cecal samples and 6 in the fecal samples, see **Table 2**). Of the 17 significant species, 8 were found at reduced abundance in 5xfAD mice relative to the WT controls (*Turicibacter* sp. H121, *Romboutsia ilealis*, *Burkholderiales* bacterium YL45, *Adlercreutzia equolifaciens*, *Duncaniella* sp. PC9, *Duncaniella* sp. B8, *Muribaculum intestinale*, and *Lactobacillus johnsonii*) and 9 were found at an increased abundance (*Lactobacillus* sp. P38, *Ligilactobacillus murinus*, *Ligilactobacillus animalis*, and six *Alistipes* species). It is important to note that closely related species (*e.g.*, a subset of the *Alistipes* species) may have abundances that are artificially over-correlated by our method for assigning BLAST hits (see **Methods**).

These findings are consistent with those in previous studies of the fecal microbiome of people with AD. Using 16S rRNA sequencing, Vogt *et. al.* reported significantly decreased levels of 7 genera of bacteria, including *Turicibacter* and *Adlercreutzia*, in people with AD relative to healthy individuals.^23^ This same study reported 7 genera of increased abundance in people with AD, including *Alistipes*, matching our observations in the cecal samples from 5xfAD mice.

### Turicibacter *spp. and Serotonin*

The finding of reduced *Turicibacter* spp. is also noteworthy because members of this genus can both metabolize and induce production of serotonin in the gut. The association between serotonin and *T. sanguinis* has been shown to be both causative and bidirectional.^39^ Along with depleted *Turicibacter*,^23^ both blood and central nervous system derived serotonin are depleted in humans with AD pathology.^40,48,49^

Here, we found three *Turicibacter* spp. – *T.* sp. H121, *T.* sp. HGF1, and *T. sanguinis* – to be significantly less abundant in 5xfAD mice (relative to WT) at 18 months of age (**Figure 4**). Our measurement of serotonin in the blood plasma of 12 and 18 month old mice showed the neurotransmitter to be at a significantly higher concentration in 18 month 5xfAD animals relative to age matched WT controls (**Figures 5d** and **S8**). Although surprising, this finding has been reported by others for 5xfAD mice.^19^ The other remarkable finding with regards to serotonin was the dramatic drop in concentration between 12 and 18 months of age, which may have larger implications for the health of the older mice. Because of the potential importance of serotonin to AD, it must be investigated whether this discrepancy between patients with AD and 5xfAD mice extends to other animal models.

Serotonin was not detected in our fecal metabolomics data; however we did detect tryptophan, which is the primary biochemical precursor of serotonin synthesis, and one of its bioactive metabolites, 5-hydroxyindoleacetic acid (5-HIAA) (**Figure S11**). 5-HIAA was significantly more abundant in 5xfAD mice relative to WT at both 8 and 12 months of age, while differences in tryptophan abundance were not significant.

It is important to note that the relationship between gut, blood, and brain serotonin is complicated and causative associations are difficult to establish. For example, although more than 90% of the body’s serotonin is made in the gut, it does not cross the blood-brain barrier.^49^

### Plasma Metabolomics

The two plasma metabolites found by LME to be significantly lower in concentration in 18 month 5xfAD animals (carnosine and lysoPC a C18:1; **Figure 5b–c**) have both been associated with AD. Among other positive effects,^50^ carnosine (a dipeptide comprising beta-alanine and histidine) and related compounds have been found to suppress beta-amyloid toxicity,^51^ and lower concentrations in the blood stream may result in a reduced capacity to mitigate AD symptoms. Interestingly, lysoPC a C18:2 (differing from lysoPC a C18:1 only by the double bond position) was previously found by Fiandaca *et. al.* to be lower in people at risk for developing AD.^32^ Several other P180 biomarkers highlighted by both Fiandaca *et. al.*^32^ and Mapstone *et. al.*^31^ showed trends in our data similar to those in people with AD, however – following quality control – they lacked adequate statistical power to provide meaningful context to this report.

### Limitations and Weaknesses

The fact that most of the variance in our data was attributable to cage effects undoubtedly obscures other interesting trends in the microbiome and metabolome, including differences due to sex, age, and genotype. The cage effects were also inexorably linked to many other factors, including maternal identity, as same-sex littermates were housed together. This linkage may cause even more pronounced cage effects, partially explaining the large proportion of variance attributable to housing ID. Interestingly, the proportion of variance attributable to housing was approximately the same for both fecal and cecal samples (**Table 2**). As we did in this work, future studies should co-house mice of the WT and study group to avoid erroneous conclusions about differences in microbiome composition. Further innovation in animal husbandry strategies may also be warranted.^52^

Our results from the plasma metabolomics, although powerful, were limited by many measurements falling outside of the quantitative range. This resulted in the necessary exclusion of many metabolites and data points and the deuteriation of statistical power for our discovery worm. Future experiments should strive to overcome this issue by using a larger sample volume, or perhaps a more sensitive metabolomics method. Depending on the question at hand, earlier timepoints (e.g., 4 months) could also be informative.

Unfortunately, we had inadequate material to perform both genomics and metabolomics on all fecal samples, forcing us to choose between the two techniques in some cases, including at the critical 18 month timepoint. We chose to prioritize genomics, and, as a result, we were only able to collect metabolomics for only a subset of the samples (namely the 4, 8, and 12 month fecal samples). Due to the advanced stage of AD pathology, it is likely that analysis of the 18-month metabolomes would have produced more meaningful results than for the younger animals.

## Conclusion

Here we describe the longitudinal gut microbiome and metabolome of a 5xfAD murine model for familial AD. We found several species of bacteria that were both over and under expressed in the AD mice relative to their WT counterparts, many of which match previous observations around the fecal microbiome in persons with AD.^23^ In line with previous studies,^19,28^ we also found large variance in the microbiome arising from housing assignment. Future studies must account for cage effects in experimental design, as, even with meticulous animal husbandry, cage effects are likely to be the largest source variance in the murine microbiome. In our case, we were able to confidently compare differences arising due to genotype among age-matched littermates of the same sex, which were housed together.

One key finding was that three species of *Turicibacter* were present at lower abundances in 18 month 5xfAD mice relative to age-matched WT controls. Some species of *Turicibacter* are known to both metabolize and regulate the production of serotonin,^39^ however serotonin concentrations were found to be significantly elevated in 18 month 5xfAD mice relative to age-matched controls. Serotonin was also observed to be nearly 10-fold less concentrated in blood plasma of 18 month mice relative to the 12 month cohort. Depending on the importance of serotonin in AD, this finding may have broad implications for the design of better murine models and our understanding of the role of the microbiome in the etiology of AD.

This research adds to our collective understanding of the microbiome gut-brain axis in neurodegeneration and serves as a foundation for further microbiome research with AD mouse models. Our findings also motivate further studies involving diet interventions, gnotobiotic and wildling mice, among other investigations.

## Supporting information

Supplemental Results

## Data Availability

Data associated with this study will be made available in appropriate public repositories upon publication and as part of the MODEL-AD consortia (https://www.model-ad.org).

## Funding and Acknowledgements

This study was funded through two supplements to the parent grant U54AG054349. SJBD is supported by the NIA Aging and Alzheimer’s Disease Training Grant (T32 AG00096-38). We couldn’t have generated the plasma metabolomics data without steadfast collaboration from Dr. Amrita Cheeta of Georgetown. The fecal and cecal metabolomes were generated at the West Coast Metabolomics Center thanks to Dr. Oliver Fiehn. Thank you to Dr. Andrea Tenner for inspiring us to examine the microbiome in this context, Dr. Shimako Kawauchi for her work developing the animal models, along with Dr. Grant MacGregor and Dr. Kim Green and the entire Model-AD team for being wonderful colleagues. We also thank Dr. Veronica Peterson for her scientific contributions, and Dr. Heather Maughan for her assistance with scientific writing.

## References

[1] Serrano-Pozo A, Frosch MP, Masliah E, Hyman BT. Neuropathological alterations in Alzheimer disease. Cold Spring Harb Perspect Med. 2011 Sep;1(1):a006189. doi: 10.1101/cshperspect.a006189. PMID: 22229116; PMCID: PMC3234452.

[2] Bellenguez C, Grenier-Boley B, Lambert JC. Genetics of Alzheimer’s disease: where we are, and where we are going. Curr Opin Neurobiol. 2020 Apr;61:40–48. doi: 10.1016/j.conb.2019.11.024. Epub 2019 Dec 18. PMID: 31863938.

[3] Rahman MA, Rahman MS, Uddin MJ, Mamum-Or-Rashid ANM, Pang MG, Rhim H. Emerging risk of environmental factors: insight mechanisms of Alzheimer’s diseases. Environ Sci Pollut Res Int. 2020 Dec;27(36):44659–44672. doi: 10.1007/s11356-020-08243-z. Epub 2020 Mar 23. PMID: 32201908.

[4] fda.gov https://www.fda.gov/drugs/news-events-human-drugs/fdas-decision-approve-new-treatment-alzheimers-disease

[5] Sevigny J, Chiao P, Bussière T, Weinreb PH, Williams L, Maier M, Dunstan R, Salloway S, Chen T, Ling Y, O’Gorman J, Qian F, Arastu M, Li M, Chollate S, Brennan MS, Quintero-Monzon O, Scannevin RH, Arnold HM, Engber T, Rhodes K, Ferrero J, Hang Y, Mikulskis A, Grimm J, Hock C, Nitsch RM, Sandrock A. The antibody aducanumab reduces Aβ plaques in Alzheimer’s disease. Nature. 2016 Sep 1;537(7618):50–6. doi: 10.1038/nature19323. Update in: Nature. 2017 Jun 21;546(7659):564. PMID: 27582220.

[6] Karlawish J, Grill JD. The approval of Aduhelm risks eroding public trust in Alzheimer research and the FDA. Nat Rev Neurol. 2021 Sep;17(9):523–524. doi: 10.1038/s41582-021-00540-6. PMID: 34267383.

[7] Knopman DS, Jones DT, Greicius MD. Failure to demonstrate efficacy of aducanumab: An analysis of the EMERGE and ENGAGE trials as reported by Biogen, December 2019. Alzheimers Dement. 2021 Apr;17(4):696–701. doi: 10.1002/alz.12213. Epub 2020 Nov 1. PMID: 33135381.

[8] Jankowsky JL, Zheng H. Practical considerations for choosing a mouse model of Alzheimer’s disease. Mol Neurodegener. 2017 Dec 22;12(1):89. doi: 10.1186/s13024-017-0231-7. PMID: 29273078; PMCID: PMC5741956.

[9] Mckean NE, Handley RR, Snell RG. A Review of the Current Mammalian Models of Alzheimer’s Disease and Challenges That Need to Be Overcome. Int J Mol Sci. 2021 Dec 6;22(23):13168. doi: 10.3390/ijms222313168. PMID: 34884970; PMCID: PMC8658123.

[10] Rabinovici GD. Late-onset Alzheimer Disease. Continuum (Minneap Minn). 2019 Feb;25(1):14–33. doi: 10.1212/CON.0000000000000700. PMID: 30707185; PMCID: PMC6548536.

[11] Baglietto-Vargas D, Forner S, Cai L, Martini AC, Trujillo-Estrada L, Swarup V, Nguyen MMT, Do Huynh K, Javonillo DI, Tran KM, Phan J, Jiang S, Kramár EA, Nuñez-Diaz C, Balderrama-Gutierrez G, Garcia F, Childs J, Rodriguez-Ortiz CJ, Garcia-Leon JA, Kitazawa M, Shahnawaz M, Matheos DP, Ma X, Da Cunha C, Walls KC, Ager RR, Soto C, Gutierrez A, Moreno-Gonzalez I, Mortazavi A, Tenner AJ, MacGregor GR, Wood M, Green KN, LaFerla FM. Generation of a humanized Aβ expressing mouse demonstrating aspects of Alzheimer’s disease-like pathology. Nat Commun. 2021 Apr 23;12(1):2421. doi: 10.1038/s41467-021-22624-z. PMID: 33893290; PMCID: PMC8065162.

[12] Bhattacharjee S, Lukiw W. Alzheimer’s disease and the microbiome. Front. Cell. Neurosci., 17 September 2013. https://doi.org/10.3389/fncel.2013.00153

[13] Jiang C, Li G, Huang P, Liu Z, Zhao B. The Gut Microbiota and Alzheimer’s Disease. J Alzheimers Dis. 2017;58(1):1–15. doi: 10.3233/JAD-161141. PMID: 28372330.

[14] Angelucci F, Cechova K, Amlerova J, Hort J. Antibiotics, gut microbiota, and Alzheimer’s disease. J Neuroinflammation. 2019 May 22;16(1):108. doi: 10.1186/s12974-019-1494-4. PMID: 31118068; PMCID: PMC6530014.

[15] Doifode T, Giridharan VV, Generoso JS, Bhatti G, Collodel A, Schulz PE, Forlenza OV, Barichello T. The impact of the microbiota-gut-brain axis on Alzheimer’s disease pathophysiology. Pharmacol Res. 2021 Feb;164:105314. doi: 10.1016/j.phrs.2020.105314. Epub 2020 Nov 25. PMID: 33246175.

[16] Kesika P, Suganthy N, Sivamaruthi BS, Chaiyasut C. Role of gut-brain axis, gut microbial composition, and probiotic intervention in Alzheimer’s disease. Life Sci. 2021 Jan 1;264:118627. doi: 10.1016/j.lfs.2020.118627. Epub 2020 Oct 22. PMID: 33169684.

[17] Heneka MT, Carson MJ, El Khoury J, Landreth GE, Brosseron F, Feinstein DL, Jacobs AH, Wyss-Coray T, Vitorica J, Ransohoff RM, Herrup K, Frautschy SA, Finsen B, Brown GC, Verkhratsky A, Yamanaka K, Koistinaho J, Latz E, Halle A, Petzold GC, Town T, Morgan D, Shinohara ML, Perry VH, Holmes C, Bazan NG, Brooks DJ, Hunot S, Joseph B, Deigendesch N, Garaschuk O, Boddeke E, Dinarello CA, Breitner JC, Cole GM, Golenbock DT, Kummer MP. Neuroinflammation in Alzheimer’s disease. Lancet Neurol. 2015 Apr;14(4):388–405. doi: 10.1016/S1474-4422(15)70016-5. PMID: 25792098; PMCID: PMC5909703.

[18] Calsolaro V, Edison P. Neuroinflammation in Alzheimer’s disease: Current evidence and future directions. Alzheimers Dement. 2016 Jun;12(6):719–32. doi: 10.1016/j.jalz.2016.02.010. Epub 2016 May 11. PMID: 27179961.

[19] Wang X, Sun G, Feng T, Zhang J, Huang X, Wang T, Xie Z, Chu X, Yang J, Wang H, Chang S, Gong Y, Ruan L, Zhang G, Yan S, Lian W, Du C, Yang D, Zhang Q, Lin F, Liu J, Zhang H, Ge C, Xiao S, Ding J, Geng M. Sodium oligomannate therapeutically remodels gut microbiota and suppresses gut bacterial amino acids-shaped neuroinflammation to inhibit Alzheimer’s disease progression. Cell Res. 2019 Oct;29(10):787–803. doi: 10.1038/s41422-019-0216-x. Epub 2019 Sep 6. PMID: 31488882; PMCID: PMC6796854.

[20] Dodiya HB, Kuntz T, Shaik SM, Baufeld C, Leibowitz J, Zhang X, Gottel N, Zhang X, Butovsky O, Gilbert JA, Sisodia SS. Sex-specific effects of microbiome perturbations on cerebral Aβ amyloidosis and microglia phenotypes. J Exp Med. 2019 Jul 1;216(7):1542–1560. doi: 10.1084/jem.20182386. Epub 2019 May 16. PMID: 31097468; PMCID: PMC6605759.

[21] Guilherme MDS, Nguyen VTT, Reinhardt C, Endres K. Impact of Gut Microbiome Manipulation in 5xFAD Mice on Alzheimer’s Disease-Like Pathology. Microorganisms. 2021 Apr 13;9(4):815. doi: 10.3390/microorganisms9040815. PMID: 33924322; PMCID: PMC8069338.

[22] Harach T, Marungruang N, Duthilleul N, Cheatham V, Mc Coy KD, Frisoni G, Neher JJ, Fåk F, Jucker M, Lasser T, Bolmont T. Reduction of Abeta amyloid pathology in APPPS1 transgenic mice in the absence of gut microbiota. Sci Rep. 2017 Feb 8;7:41802. doi: 10.1038/srep41802. Erratum in: Sci Rep. 2017 Jul 10;7:46856. PMID: 28176819; PMCID: PMC5297247.

[23] Vogt NM, Kerby RL, Dill-McFarland KA, Harding SJ, Merluzzi AP, Johnson SC, Carlsson CM, Asthana S, Zetterberg H, Blennow K, Bendlin BB, Rey FE. Gut microbiome alterations in Alzheimer’s disease. Sci Rep. 2017 Oct 19;7(1):13537. doi: 10.1038/s41598-017-13601-y. PMID: 29051531; PMCID: PMC5648830.

[24] Haran JP, Bhattarai SK, Foley SE, Dutta P, Ward DV, Bucci V, McCormick BA. Alzheimer’s Disease Microbiome Is Associated with Dysregulation of the Anti-Inflammatory P-Glycoprotein Pathway. mBio. 2019 May 7;10(3):e00632–19. doi: 10.1128/mBio.00632-19. PMID: 31064831; PMCID: PMC6509190.

[25] MahmoudianDehkordi S, Arnold M, Nho K, Ahmad S, Jia W, Xie G, Louie G, Kueider-Paisley A, Moseley MA, Thompson JW, St John Williams L, Tenenbaum JD, Blach C, Baillie R, Han X, Bhattacharyya S, Toledo JB, Schafferer S, Klein S, Koal T, Risacher SL, Kling MA, Motsinger-Reif A, Rotroff DM, Jack J, Hankemeier T, Bennett DA, De Jager PL, Trojanowski JQ, Shaw LM, Weiner MW, Doraiswamy PM, van Duijn CM, Saykin AJ, Kastenmüller G, Kaddurah-Daouk R; Alzheimer’s Disease Neuroimaging Initiative and the Alzheimer Disease Metabolomics Consortium. Altered bile acid profile associates with cognitive impairment in Alzheimer’s disease-An emerging role for gut microbiome. Alzheimers Dement. 2019 Jan;15(1):76–92. doi: 10.1016/j.jalz.2018.07.217. Epub 2018 Oct 15. Erratum in: Alzheimers Dement. 2019 Apr;15(4):604. PMID: 30337151; PMCID: PMC6487485.

[26] Oakley H, Cole SL, Logan S, Maus E, Shao P, Craft J, Guillozet-Bongaarts A, Ohno M, Disterhoft J, Van Eldik L, Berry R, Vassar R. Intraneuronal beta-amyloid aggregates, neurodegeneration, and neuron loss in transgenic mice with five familial Alzheimer’s disease mutations: potential factors in amyloid plaque formation. J Neurosci. 2006 Oct 4;26(40):10129–40. doi: 10.1523/JNEUROSCI.1202-06.2006. PMID: 17021169; PMCID: PMC6674618.

[27] Forner S, Kawauchi S, Balderrama-Gutierrez G, Kramár EA, Matheos DP, Phan J, Javonillo DI, Tran KM, Hingco E, da Cunha C, Rezaie N, Alcantara JA, Baglietto-Vargas D, Jansen C, Neumann J, Wood MA, MacGregor GR, Mortazavi A, Tenner AJ, LaFerla FM, Green KN. Systematic phenotyping and characterization of the 5xFAD mouse model of Alzheimer’s disease. Sci Data. 2021 Oct 15;8(1):270. doi: 10.1038/s41597-021-01054-y. PMID: 34654824; PMCID: PMC8519958.

[28] Oblak AL, Lin PB, Kotredes KP, Pandey RS, Garceau D, Williams HM, Uyar A, O’Rourke R, O’Rourke S, Ingraham C, Bednarczyk D, Belanger M, Cope ZA, Little GJ, Williams SG, Ash C, Bleckert A, Ragan T, Logsdon BA, Mangravite LM, Sukoff Rizzo SJ, Territo PR, Carter GW, Howell GR, Sasner M, Lamb BT. Comprehensive Evaluation of the 5XFAD Mouse Model for Preclinical Testing Applications: A MODEL-AD Study. Front Aging Neurosci. 2021 Jul 23;13:713726. doi: 10.3389/fnagi.2021.713726. PMID: 34366832; PMCID: PMC8346252.

[29] Weihe C, Avelar-Barragan J. Next generation shotgun library preparation for Illumina sequencing – low volume. protocols.io. 2021 Jul. doi: 10.17504/protocols.io.bvv8n69w

[30] Zheng J, Wittouck S, Salvetti E, Franz CMAP, Harris HMB, Mattarelli P, O’Toole PW, Pot B, Vandamme P, Walter J, Watanabe K, Wuyts S, Felis GE, Gänzle MG, Lebeer S. A taxonomic note on the genus Lactobacillus: Description of 23 novel genera, emended description of the genus Lactobacillus Beijerinck 1901, and union of Lactobacillaceae and Leuconostocaceae. Int J Syst Evol Microbiol. 2020 Apr;70(4):2782–2858. doi: 10.1099/ijsem.0.004107. Epub 2020 Apr 15. PMID: 32293557.

[31] Mapstone M, Cheema AK, Fiandaca MS, Zhong X, Mhyre TR, MacArthur LH, Hall WJ, Fisher SG, Peterson DR, Haley JM, Nazar MD, Rich SA, Berlau DJ, Peltz CB, Tan MT, Kawas CH, Federoff HJ. Plasma phospholipids identify antecedent memory impairment in older adults. Nat Med. 2014 Apr;20(4):415–8. doi: 10.1038/nm.3466. Epub 2014 Mar 9. PMID: 24608097; PMCID: PMC5360460.

[32] Fiandaca MS, Zhong X, Cheema AK, Orquiza MH, Chidambaram S, Tan MT, Gresenz CR, FitzGerald KT, Nalls MA, Singleton AB, Mapstone M, Federoff HJ. Plasma 24-metabolite Panel Predicts Preclinical Transition to Clinical Stages of Alzheimer’s Disease. Front Neurol. 2015 Nov 12;6:237. doi: 10.3389/fneur.2015.00237. PMID: 26617567; PMCID: PMC4642213.

[33] Mapstone M, Lin F, Nalls MA, Cheema AK, Singleton AB, Fiandaca MS, Federoff HJ. What success can teach us about failure: the plasma metabolome of older adults with superior memory and lessons for Alzheimer’s disease. Neurobiology of aging. 2016. doi: 10.1016/j.neurobiolaging.2016.11.007. PubMed PMID: 27939698.

[34] Zhang Z, Ersoz E, Lai C-Q, Todhunter RJ, Tiwari HK, Gore M, Bradbury PJ, Yu J, Arnett DK, Ordovas JM, Buckler ES, Cho RJ, Mindrinos M, Richards DR, Sapolsky RJ, Anderson M, Drenkard E, Dewdney J, Reuber TL, Stammers M, Federspiel N, Theologis A, Yang WH, Hubbell E, Au M, Chung EY, Lashkari D, Lemieux B, Dean C, Lipshutz RJ, Ausubel FM, Davis RW, Oefner PJ, Bradbury PJ, Zhang Z, Kroon DE, Casstevens TM, Ramdoss Y, Buckler ES, Glaubitz JC, Casstevens TM, Lu F, Harriman J, Elshire RJ, Sun Q, Buckler ES, Lenné JM, Takan JP, Mgonja MA, Manyasa EO, Kaloki P, Wanyera N, et al. 2014. R: a language and environment for statistical computing. R Foundation for Statistical Computing, Vienna, Austria. http://www.R-project.org/.

[35] Wickham H. 2009. ggplot2: elegant graphics for data analysis. Springer-Verlag, New York, NY

[36] Kolde R. 2019. pheatmap: pretty heatmaps. https://cran.r-project.org/web/packages/pheatmap/index.html.

[37] Oksanen J, Blanchet FG, Friendly M, Kindt R, Legendre P, McGlinn D, Minchin PR, O’Hara RB, Simpson GL, Solymos P, Stevens MHH, Szoecs E, Wagner H. 2017 vegan: community ecology package. https://github.com/vegandevs/vegan. Accessed on 1 April 2021

[38] Kuznetsova, A., Brockhoff, P. B., & Christensen, R. H. B. (2017). {lmerTest} Package: Tests in Linear Mixed Effects Models. Journal of Statistical Software, 82(13), 1–26. https://doi.org/10.18637/jss.v082.i13

[39] Fung TC, Vuong HE, Luna CDG, Pronovost GN, Aleksandrova AA, Riley NG, Vavilina A, McGinn J, Rendon T, Forrest LR, Hsiao EY. Intestinal serotonin and fluoxetine exposure modulate bacterial colonization in the gut. Nat Microbiol. 2019 Dec;4(12):2064–2073. doi: 10.1038/s41564-019-0540-4. Epub 2019 Sep 2. PMID: 31477894; PMCID: PMC6879823.

[40] Aaldijk E, Vermeiren Y. The role of serotonin within the microbiota-gut-brain axis in the development of Alzheimer’s disease: A narrative review. Ageing Res Rev. 2022 Mar;75:101556. doi: 10.1016/j.arr.2021.101556. Epub 2022 Jan 3. PMID: 34990844.

[41] de la Cuesta-Zuluaga J, Kelley ST, Chen Y, Escobar JS, Mueller NT, Ley RE, McDonald D, Huang S, Swafford AD, Knight R, Thackray VG. Age- and Sex-Dependent Patterns of Gut Microbial Diversity in Human Adults. mSystems. 2019 May 14;4(4):e00261-19. doi: 10.1128/mSystems.00261-19. PMID: 31098397; PMCID: PMC6517691.

[42] Yun Y, Kim HN, Kim SE, Heo SG, Chang Y, Ryu S, Shin H, Kim HL. Comparative analysis of gut microbiota associated with body mass index in a large Korean cohort. BMC Microbiol. 2017 Jul 4;17(1):151. doi: 10.1186/s12866-017-1052-0. PMID: 28676106; PMCID: PMC5497371.

[43] Claesson MJ, Jeffery IB, Conde S, Power SE, O’Connor EM, Cusack S, Harris HM, Coakley M, Lakshminarayanan B, O’Sullivan O, Fitzgerald GF, Deane J, O’Connor M, Harnedy N, O’Connor K, O’Mahony D, van Sinderen D, Wallace M, Brennan L, Stanton C, Marchesi JR, Fitzgerald AP, Shanahan F, Hill C, Ross RP, O’Toole PW. Gut microbiota composition correlates with diet and health in the elderly. Nature. 2012 Aug 9;488(7410):178–84. doi: 10.1038/nature11319. PMID: 22797518.

[44] Deloris Alexander A, Orcutt RP, Henry JC, Baker J Jr, Bissahoyo AC, Threadgill DW. Quantitative PCR assays for mouse enteric flora reveal strain-dependent differences in composition that are influenced by the microenvironment. Mamm Genome. 2006 Nov;17(11):1093–104. doi: 10.1007/s00335-006-0063-1. Epub 2006 Nov 7. PMID: 17091319.

[45] Hildebrand F, Nguyen TL, Brinkman B, Yunta RG, Cauwe B, Vandenabeele P, Liston A, Raes J. Inflammation-associated enterotypes, host genotype, cage and inter-individual effects drive gut microbiota variation in common laboratory mice. Genome Biol. 2013 Jan 24;14(1):R4. doi: 10.1186/gb-2013-14-1-r4. PMID: 23347395; PMCID: PMC4053703.

[46] Ericsson AC, Gagliardi J, Bouhan D, Spollen WG, Givan SA, Franklin CL. The influence of caging, bedding, and diet on the composition of the microbiota in different regions of the mouse gut. Sci Rep. 2018 Mar 6;8(1):4065. doi: 10.1038/s41598-018-21986-7. PMID: 29511208; PMCID: PMC5840362.

[47] Singh G, Brass A, Cruickshank SM, Knight CG. Cage and maternal effects on the bacterial communities of the murine gut. Sci Rep. 2021 May 10;11(1):9841. doi: 10.1038/s41598-021-89185-5. PMID: 33972615; PMCID: PMC8110963.

[48] Tajeddinn W, Fereshtehnejad SM, Seed Ahmed M, Yoshitake T, Kehr J, Shahnaz T, Milovanovic M, Behbahani H, Höglund K, Winblad B, Cedazo-Minguez A, Jelic V, Järemo P, Aarsland D. Association of Platelet Serotonin Levels in Alzheimer’s Disease with Clinical and Cerebrospinal Fluid Markers. J Alzheimers Dis. 2016 May 4;53(2):621–30. doi: 10.3233/JAD-160022. PMID: 27163811.

[49] El-Merahbi R, Löffler M, Mayer A, Sumara G. The roles of peripheral serotonin in metabolic homeostasis. FEBS Lett. 2015 Jul 8;589(15):1728–34. doi: 10.1016/j.febslet.2015.05.054. Epub 2015 Jun 9. PMID: 26070423.

[50] Schön M, Mousa A, Berk M, Chia WL, Ukropec J, Majid A, Ukropcová B, de Courten B. The Potential of Carnosine in Brain-Related Disorders: A Comprehensive Review of Current Evidence. Nutrients. 2019 May 28;11(6):1196. doi: 10.3390/nu11061196. PMID: 31141890; PMCID: PMC6627134.

[51] Greco V, Naletova I, Ahmed IMM, Vaccaro S, Messina L, La Mendola D, Bellia F, Sciuto S, Satriano C, Rizzarelli E. Hyaluronan-carnosine conjugates inhibit Aβ aggregation and toxicity. Sci Rep. 2020 Sep 29;10(1):15998. doi: 10.1038/s41598-020-72989-2. PMID: 32994475; PMCID: PMC7524733.

[52] Robertson SJ, Lemire P, Maughan H, Goethel A, Turpin W, Bedrani L, Guttman DS, Croitoru K, Girardin SE, Philpott DJ. Comparison of Co-housing and Littermate Methods for Microbiota Standardization in Mouse Models. Cell Rep. 2019 May 7;27(6):1910–1919.e2. doi: 10.1016/j.celrep.2019.04.023. PMID: 31067473.

