## Supplemental Figures and Tables for "Longitudinal analysis of the gut microbiome in the 5xfAD mouse model of Alzheimer’s disease"

Sage J. B. Dunham*[a], Katelyn A. McNair[b], Eric D. Adams[a], Julio Avelar-Barragan[a], Stefania Forner[c], Mark Mapstone[d], Katrine L. Whiteson[a]*

[a]Department of Molecular Biology & Biochemistry
University of California Irvine
3315 McGaugh Hall
Irvine, CA, 92697

[b]Department of Computational Science
University of California Irvine
3019 Donald Bren Hall
Irvine, CA, 92697

[c]Institute for Memory Impairments and Neurological Disorders (UCI MIND)
University of California Irvine
Biological Sciences III, 2642
Irvine, CA, 92697

[d]Department of Neurology
University of California Irvine
Irvine, CA, United States

### Contents:

**Table S1:** Overview of shotgun metagenomic sequencing demographics.
**Figure S1**: Microbiome alpha diversity with males and females considered separately.
**Figure S2**: PCoA of the microbiomes with sample type and sex considered separately.
**Figure S3**: Random forest analysis of the microbiome by sample type, age, and sex.
**Figure S4**: Relative abundance plots for the 17 significant species at 18 months of age by LME.
**Figure S5**: Alpha and beta diversity metrics for plasma metabolomics.
**Figure S6**: Random forest analysis for plasma metabolomics.
**Table S2:** LME of plasma metabolomics.
**Figure S7**: Significant plasma metabolites by LME.
**Figure S8**: Serotonin at 18 months of age.
**Figure S9**: Spearman correlation between plasma metabolites and cecal microbiome.
**Figure S10**: Spearman correlation between plasma metabolites and fecal microbiome.
**Figure S11**: Normalized abundances of tryptophan and 5-HIAA in fecal samples.

**Table S1**. Overview of shotgun metagenomic sequencing demographics.

| | **Cecal Microbiome** | | **Fecal Microbiome** | | | |
|---|---|---|---|---|---|---|
| | 4 mo | 18 mo | 4 mo | 8 mo | 12 mo | 18 mo |
| **WT Samples (female\|male)** | 18 (9\|9) | 20 (15\|5) | 38 (19\|19) | 14 (7\|7) | 18 (9\|9) | 20 (15\|5) |
| **5xfAD Samples (female\|male)** | 13 (6\|7) | 15 (6\|9) | 33 (16\|17) | 13 (6\|7) | 18 (9\|9) | 16 (6\|10) |
| **Avg. Reads* (million; ± RSD)** | 1.5 (±31%) | 1.9 (± 31%) | 1.3 (± 26%) | 1.1 (± 26%) | 1.3 (± 33%) | 1.6 (± 22%) |
| **BLAST Hit Success rate** | 18% | 19% | 24% | 20% | 26% | 28% |

*after filtering to remove mouse and rat DNA

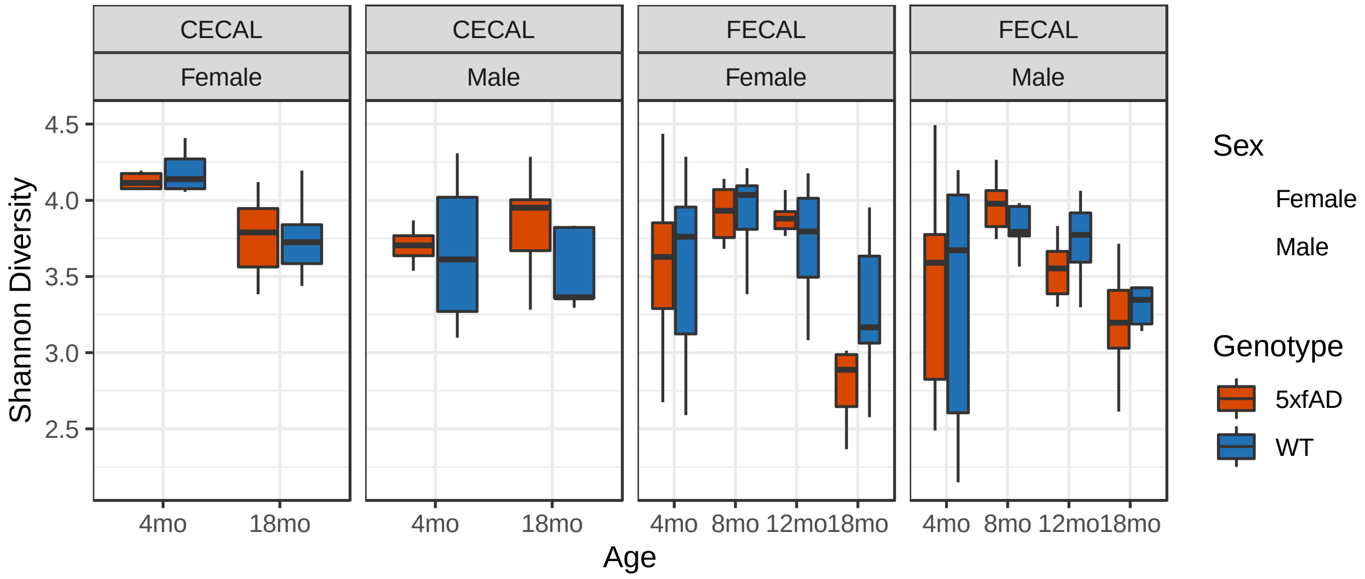


**Figure S1**. Alpha diversity of the cecal and fecal microbiomes of 5xfAD and WT mice from 4 to 18 months of age with males and females considered separately. No 5xfAD vs WT comparisons were significantly different (t-test p-value > 0.05 in all cases).

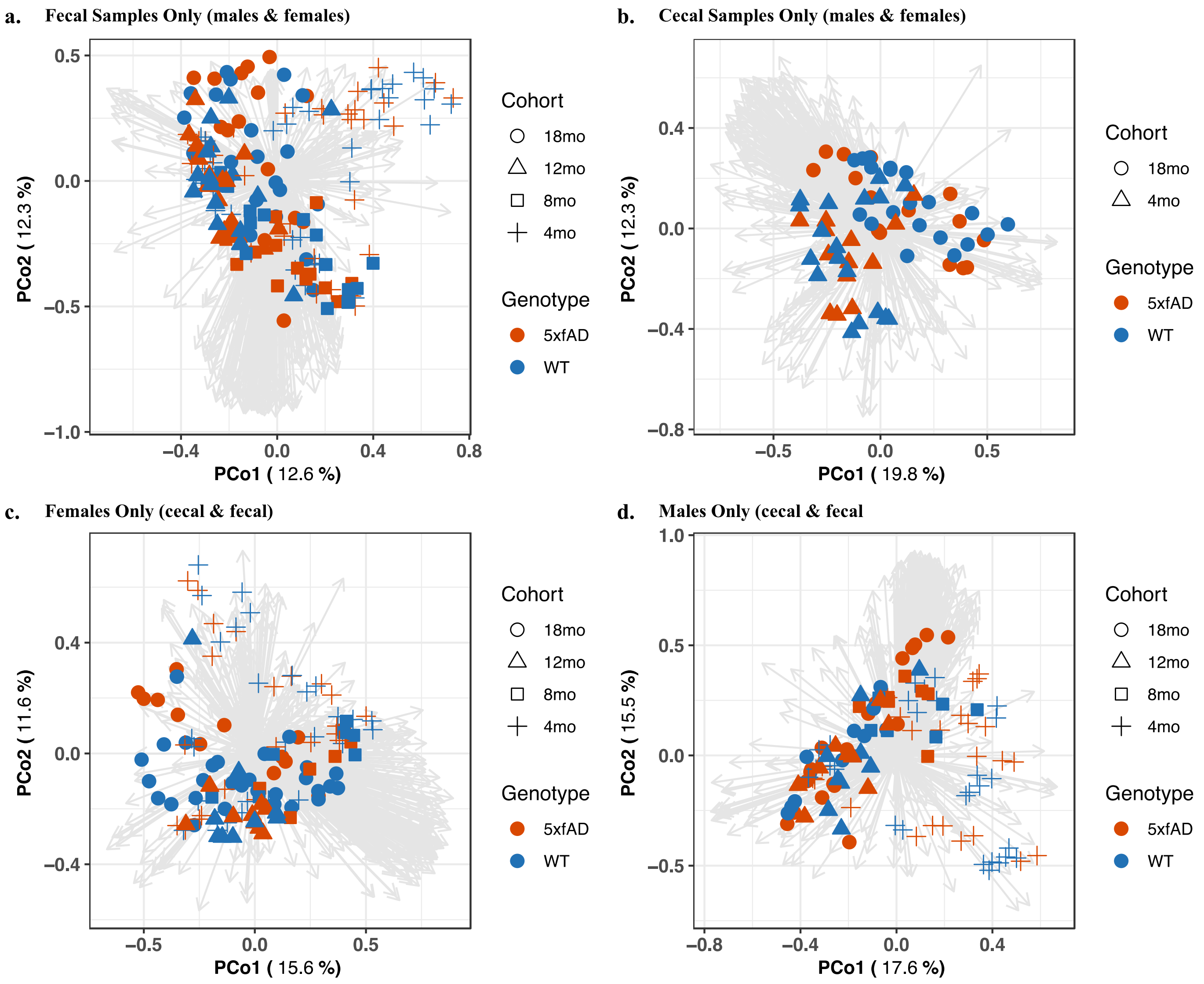


**Figure S2**. PCoA using fecal samples only (**a**), cecal samples only (**b**), females only (**c**), and males only (**d**).

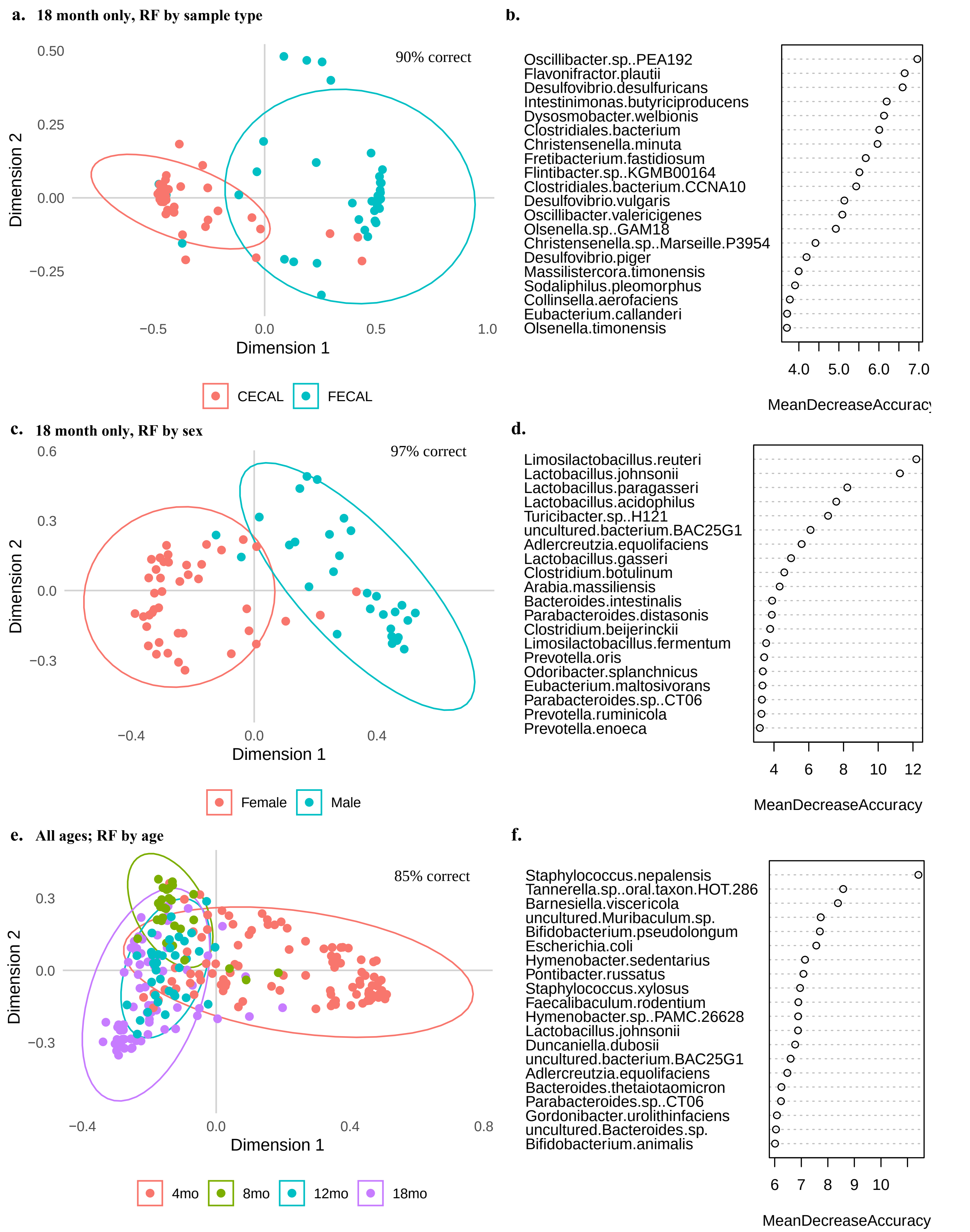


**Figure S3**. Random forest analysis of: 18 month samples with grouping by sample type (**a-b**); 18 month samples with grouping by sex (**c-d**); and all ages with grouping by age (**e-f**).

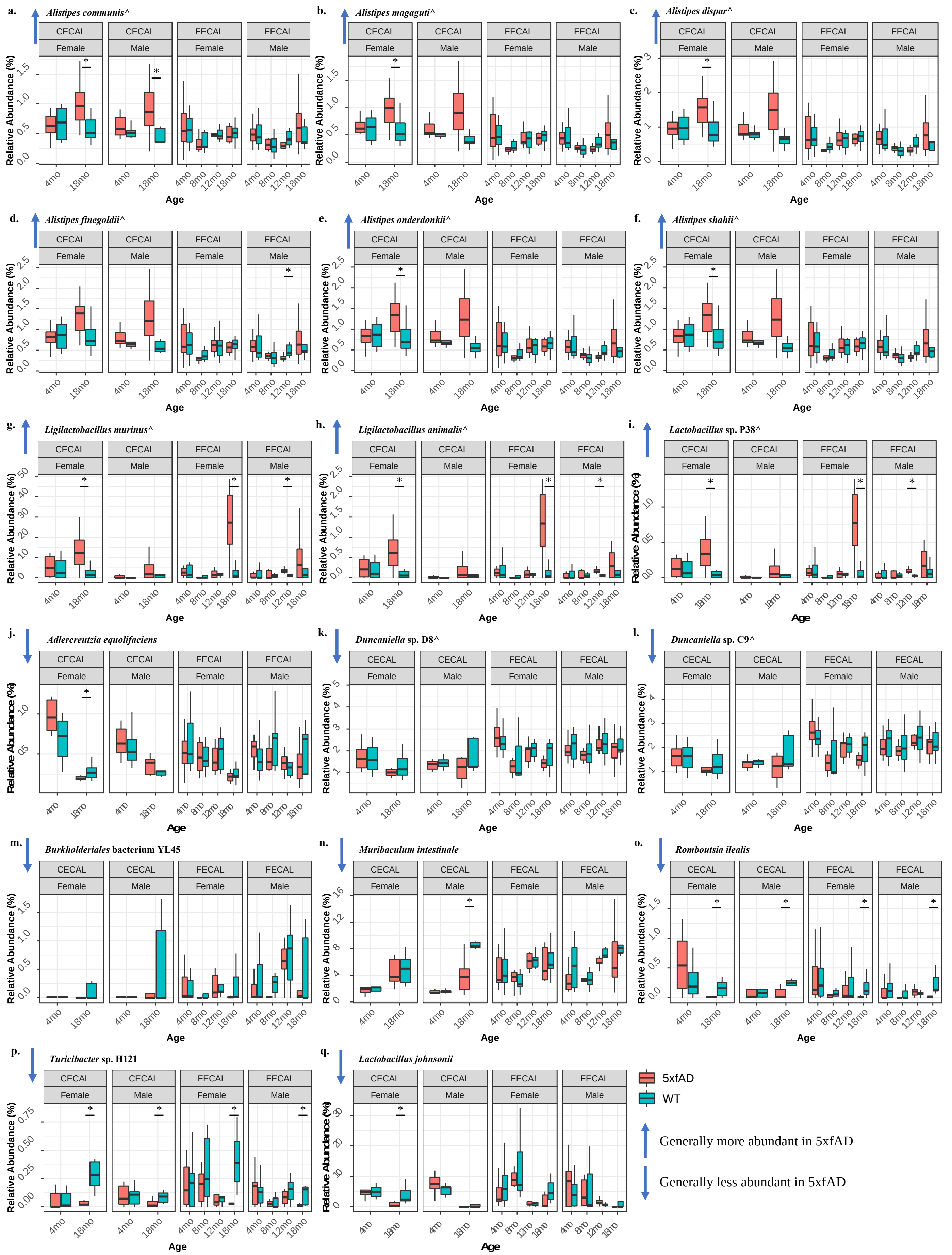


**Figure S4**. Relative abundance plots for the 17 species with significantly different abundances (padj < 0.05) by all three LME comparisons. The plots are faceted by sample type and sex. Species that were significantly different within each facet (p < 0.05 by the Wilcoxon signed rank test) are denoted with an asterisk. The box plots show the median relative abundance +/- the first and third quartiles, with the whiskers showing the range or 1.5 x the interquartile range, whichever was less.

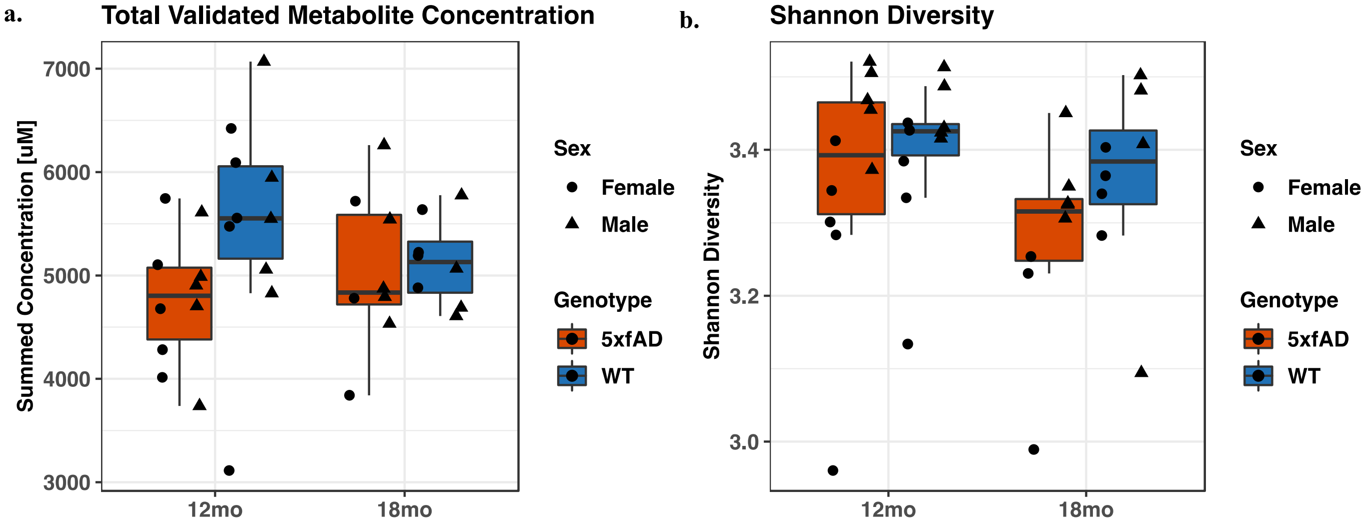


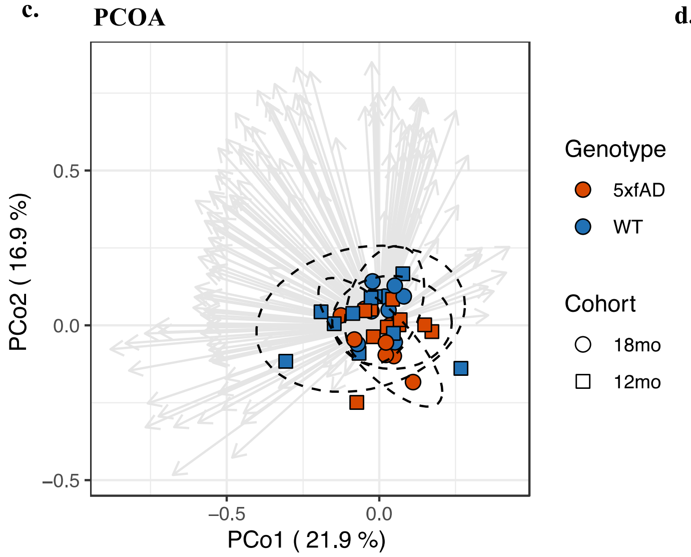


**d. PERMANOVA**

| | Df | SumsOfSqs | MeanSqs | F.Model | R2 | Pr(>F) |
|---|---|---|---|---|---|---|
| **Genotype** | 1 | 0.2769 | 0.027690 | 1.90211 | 0.04891 | 0.104 |
| **Sex** | 1 | 0.03924 | 0.039241 | 2.69553 | 0.06931 | 0.037 |
| **Age** | 1 | 0.01423 | 0.014235 | 0.97782 | 0.02514 | 0.396 |
| **Housing ID** | 14 | 0.22293 | 0.015924 | 1.09382 | 0.39377 | 0.357 |
| **Residuals** | 18 | 0.26204 | 0.014558 | | 0.46286 | |
| **Total** | 35 | 0.56614 | | | 1.00000 | |

**Figure S5**. Alpha and beta diversity metrics for plasma metabolomics: (**a**) summed concentrations for all metabolites; (**b**) Shannon diversity; (**c**) PCoA; and (**d**) PERMANOVA.

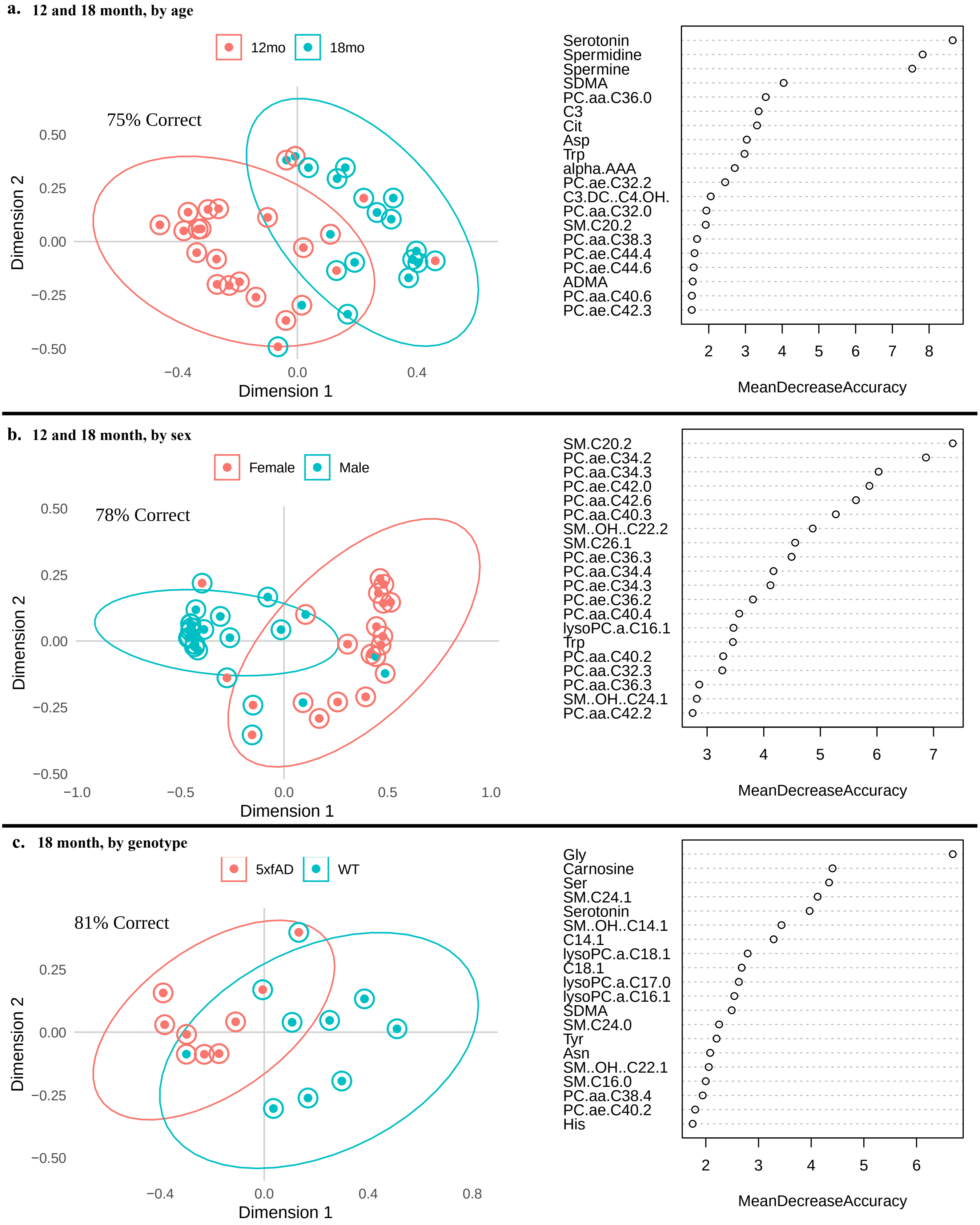


**Figure S6**. Random forest analysis of the plasma metabolome: (**a**) 12 and 18 months by age; (**b**) 12 and 18 months by sex; and (**c**) 18 month samples only by genotype.

**Table S2.** Significant plasma metabolites by LME analysis. Comparisons made between metabolomes of 12 and 18-month WT and 5XFAD animals.

| **18 month samples by genotype** | | |
|---|---|---|
| **Metabolite** | **padj** | **5xfAD vs WT** |
| Carnosine | <0.005 | Lower in 5xfAD |
| **12 & 18 month samples by genotype** | | |
| **Metabolite** | **padj** | **5xfAD vs WT** |
| Lyso PC a C18:1 | <0.05 | Lower in 5xfAD |
| Carnosine | <0.05 | - |
| **12 & 18 month samples by age** | | |
| **Metabolite** | **padj** | **18 vs 12 mo** |
| Serotonin | <0.005 | Lower in 18 mo |
| Spermine | <0.05 | Lower in 18 mo |
| Spermidine | <0.05 | Lower in 18 mo |
| Glutamic acid | <0.05 | Lower in 18 mo |
| **12 & 18 month samples by sex** | | |
| **Metabolite** | **padj** | **F vs M** |
| SM C20:2 | <0.005 | Lower in F |
| PC ae C42:0 | <0.01 | Lower in F |
| PC aa C34:3 | <0.05 | Lower in F |
| PC aa C42:6 | <0.05 | Lower in F |
| Lysine | <0.05 | Lower in F |
| SM (OH) C24:1 | <0.05 | Higher in F |
| PC aa C36:4 | <0.05 | Lower in F |
| PC aa C36:5 | <0.05 | Lower in F |
| PC ae C38:0 | <0.05 | Lower in F |
| PC aa 36:3 | <0.05 | Lower in F |
| PC aa 38:6 | <0.05 | Lower in F |

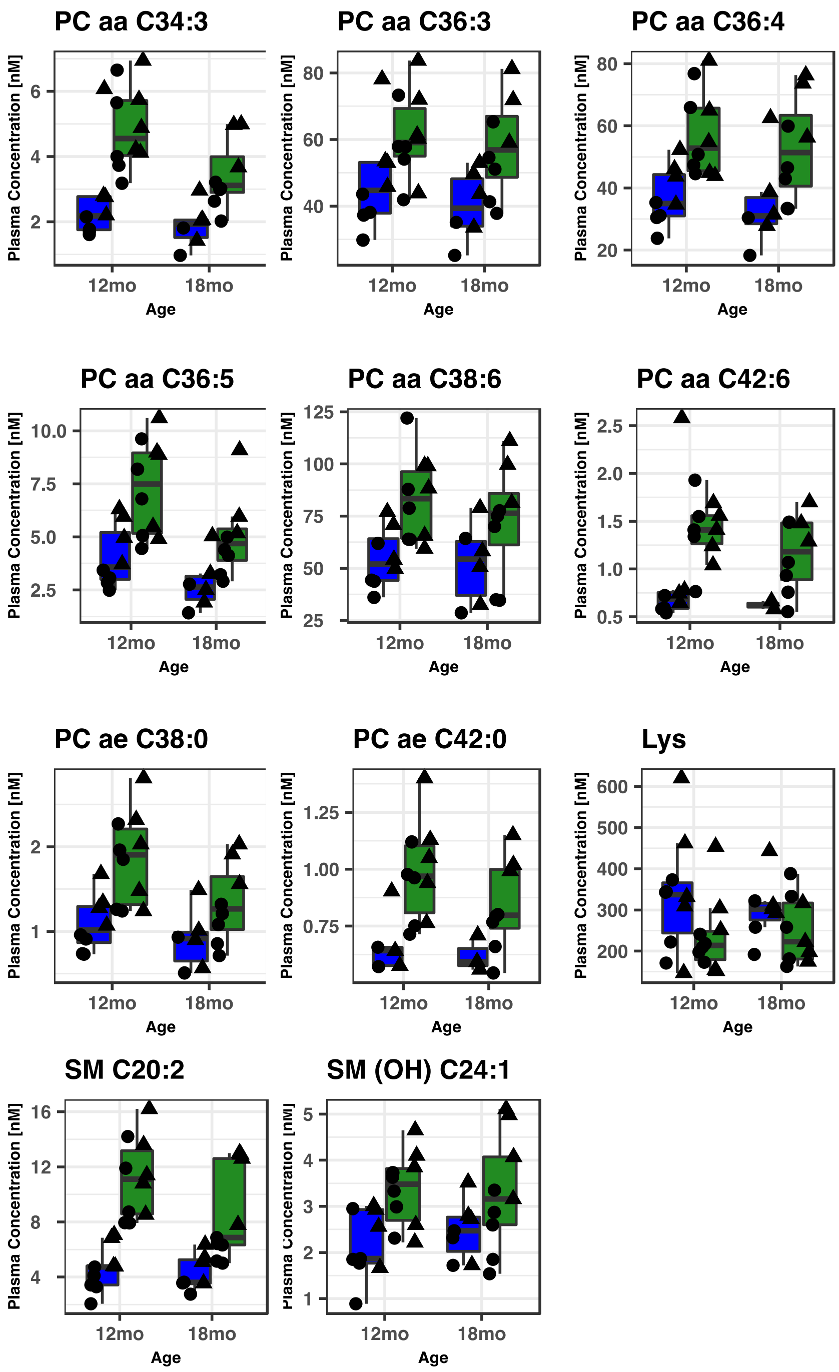


**Figure S7**. Significantly different plasma metabolites with respect to sex as revealed by LME.

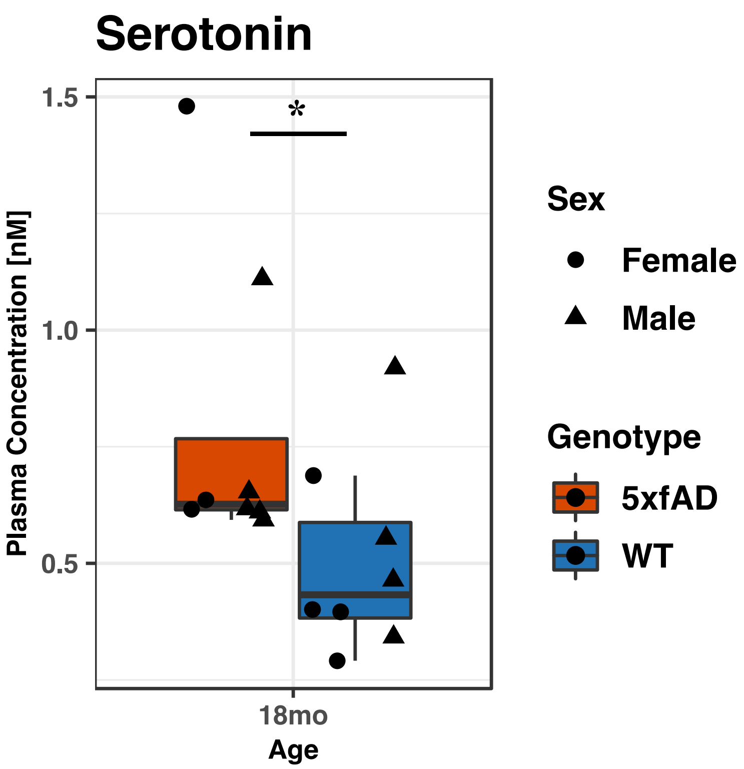


**Figure S8**. Concentration of serotonin in the plasma of 18 month 5xfAD animals.

**Figure S9**. Spearman correlation between all 137 quantified plasma metabolites and the 100 most abundant microbes in the cecal samples.

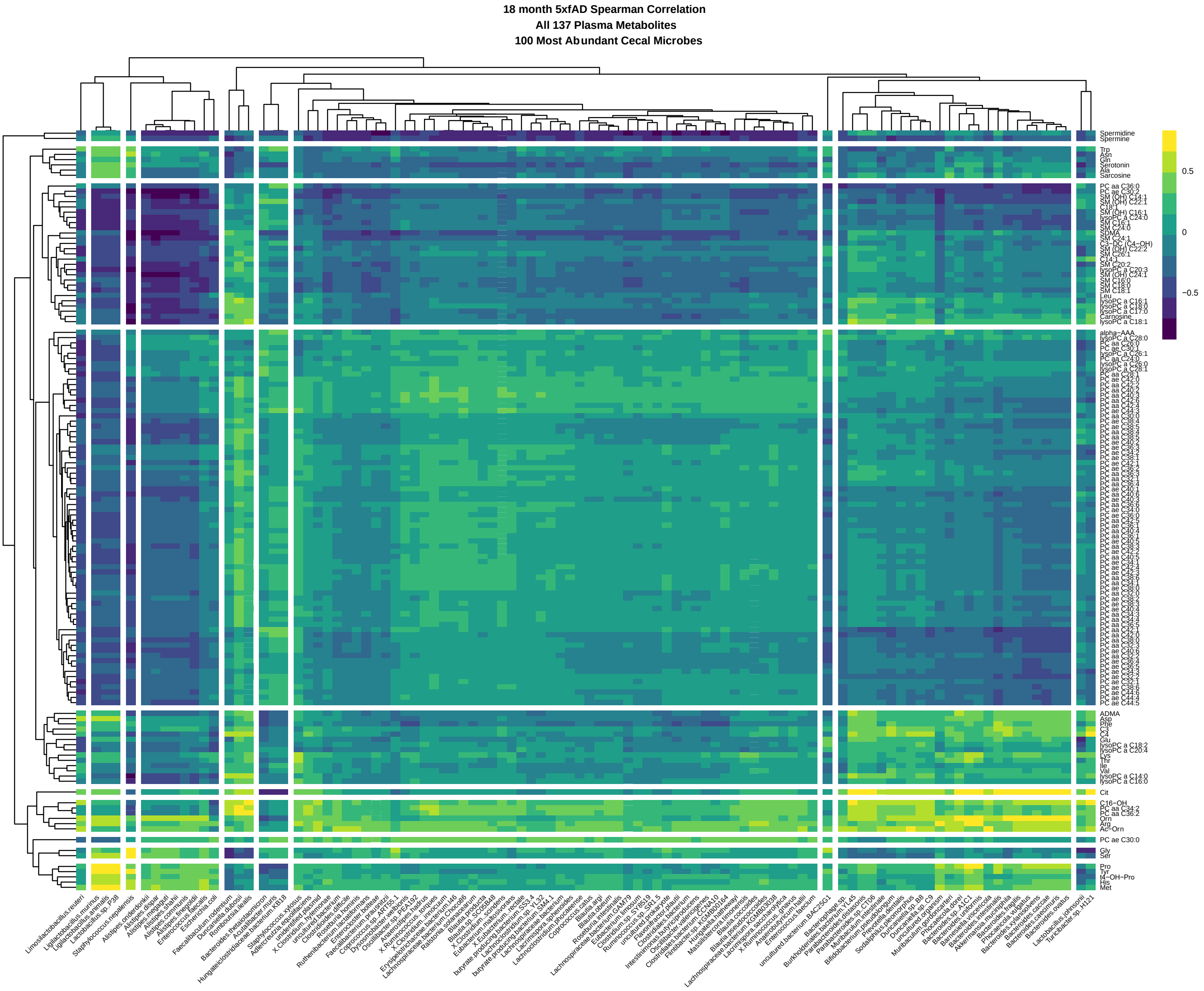

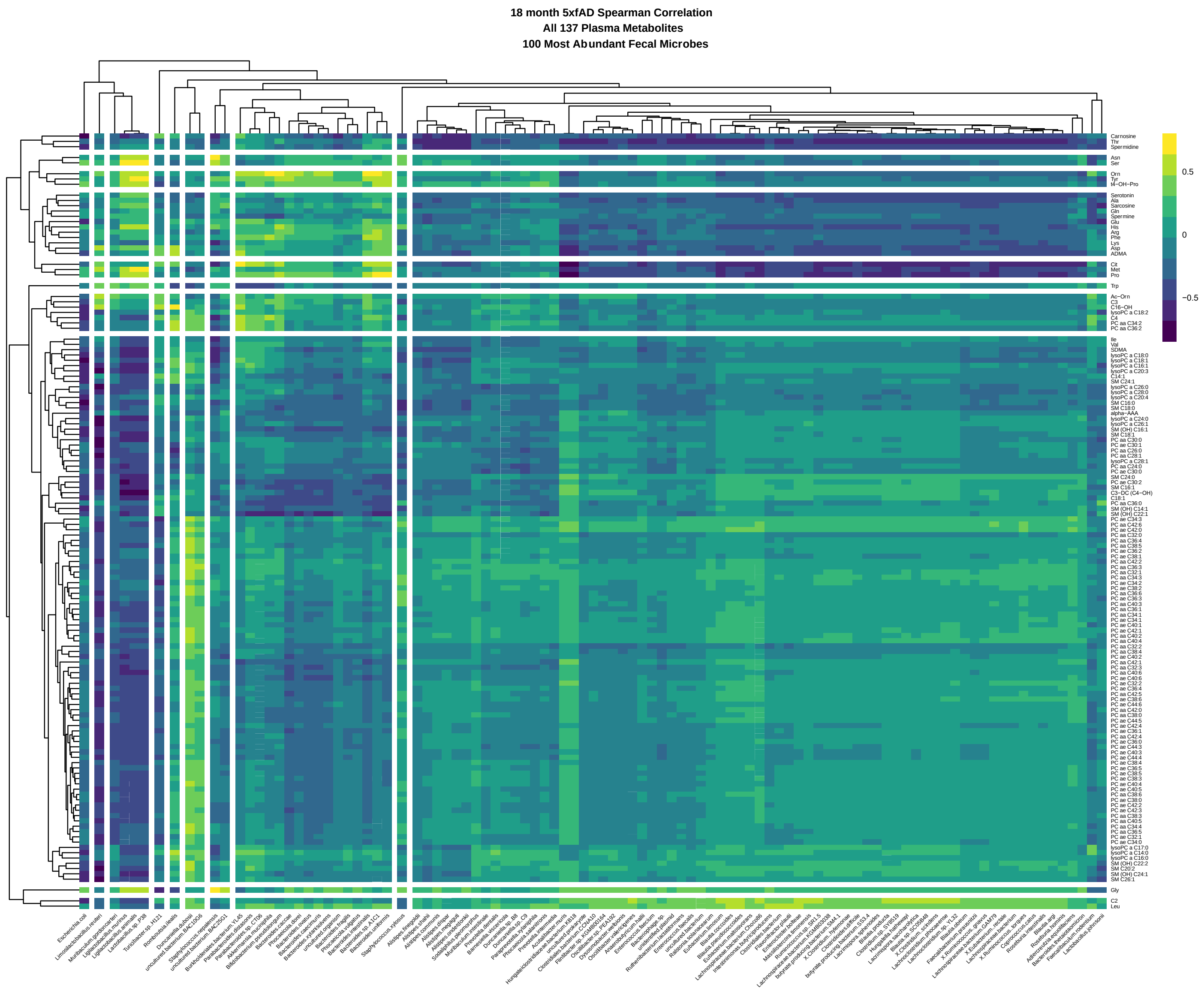


**Figure S10**. Spearman correlation between all 137 quantified plasma metabolites and the 100 most abundant microbes in the fecal samples.

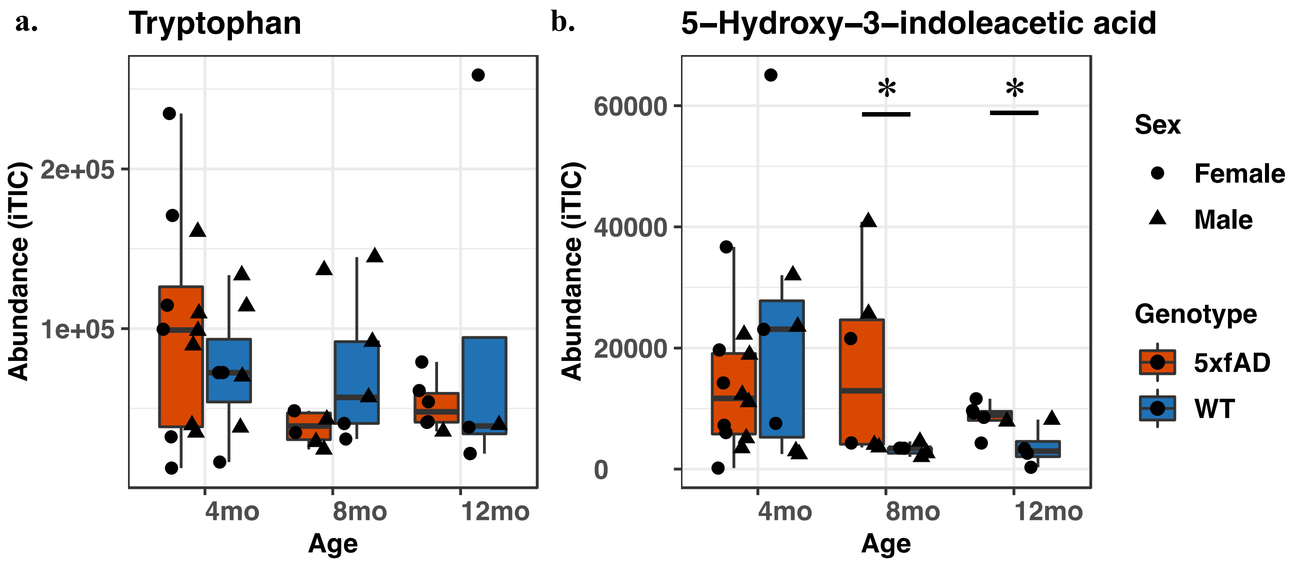


**Figure S11.** Normalized abundances of (**a**) tryptophan and (**b**) 5-hydroxyindoleacetic acid (5-HIAA) in 4, 8, and 12 month fecal samples. 5-HIAA was differentially abundant when comparing WT and 5xfAD samples at 8 and 12 months of age ($p < 0.05$ by the Wilcoxon signed rank test). All other WT vs 5xfAD comparisons were not significantly different.
