## Supplemental Results for "Longitudinal analysis of the gut microbiome in the 5xfAD mouse model of Alzheimer’s disease"

Sage J. B. Dunham*[a], Katelyn A. McNair[b], Eric D. Adams[a], Julio Avelar-Barragan[a], Stefania Forner[c], Mark Mapstone[d], Katrine L. Whiteson[a]*

[a]Department of Molecular Biology & Biochemistry
University of California Irvine
3315 McGaugh Hall
Irvine, CA, 92697

[b]Department of Computational Science
University of California Irvine
3019 Donald Bren Hall
Irvine, CA, 92697

[c]Institute for Memory Impairments and Neurological Disorders (UCI MIND)
University of California Irvine
Biological Sciences III, 2642
Irvine, CA, 92697

[d]Department of Neurology
University of California Irvine
Irvine, CA, United States

Untargeted metabolomics (via HILIC-QTOF-MS) was performed for those samples with adequate material, namely a subset of fecal samples (n = 40) from the 4, 8, and 12 month old mice. There was insufficient material to perform both metabolomics and genomics for the 18 month mice, and therefore there are no metabolomics data for this age cohort. Relative abundances were produced for 684 named metabolites with varying degrees of identification confidence, as well as more than 3500 unidentifiable ions. For this analysis we considered named metabolites only.
PERMANOVA with genotype, sex, age, and housing ID, revealed age to be the only significant parameter ($p = 0.013$), accounting for approximately 9% of the total variance (**Table SR1**). Remarkably, housing ID matched the 50% variance observed in the microbiome, however this parameter was not significant ($p = 0.34$).

| | Df | SumsOfSqs | MeanSqs | F.Model | R2 | Pr(>F) |
|---|---|---|---|---|---|---|
| **Genotype** | 1 | 0.02859 | 0.028589 | 0.53664 | 0.01311 | 0.896 |
| **Age** | 2 | 0.18891 | 0.094457 | 1.77303 | 0.08664 | 0.037 |
| **Sex** | 1 | 0.04133 | 0.041326 | 0.77571 | 0.01895 | 0.645 |
| **Housing ID** | 19 | 1.06923 | 0.056275 | 1.05633 | 0.49037 | 0.343 |
| **Residuals** | 16 | 0.85239 | 0.053274 | | 0.39092 | |
| **Total** | 39 | 2.18045 | | | 1.00000 | |

**Table SR1**. Permutational multivariate analysis of variance (PERMANOVA) of the model animal fecal metabolomes.

LME was performed to find metabolites whose abundances significantly differed between the WT and 5xfAD mice, including WT vs 5xfAD for each age and sex individually and WT vs 5xfAD for the full data set. The 12 month females were the only samples with significantly different metabolites (padj < 0.05). All significant metabolites (nialamide, fluorene, the tripeptide Ala-Pro-Lys, ferulic acid, 4 acetylbutyric acid, and disaccharide 16) were more abundant in 12 month 5xfAD female samples relative to the age and sex matched controls (see **Figure SR1**). No significant metabolites were found with other WT vs 5xfAD comparisons.

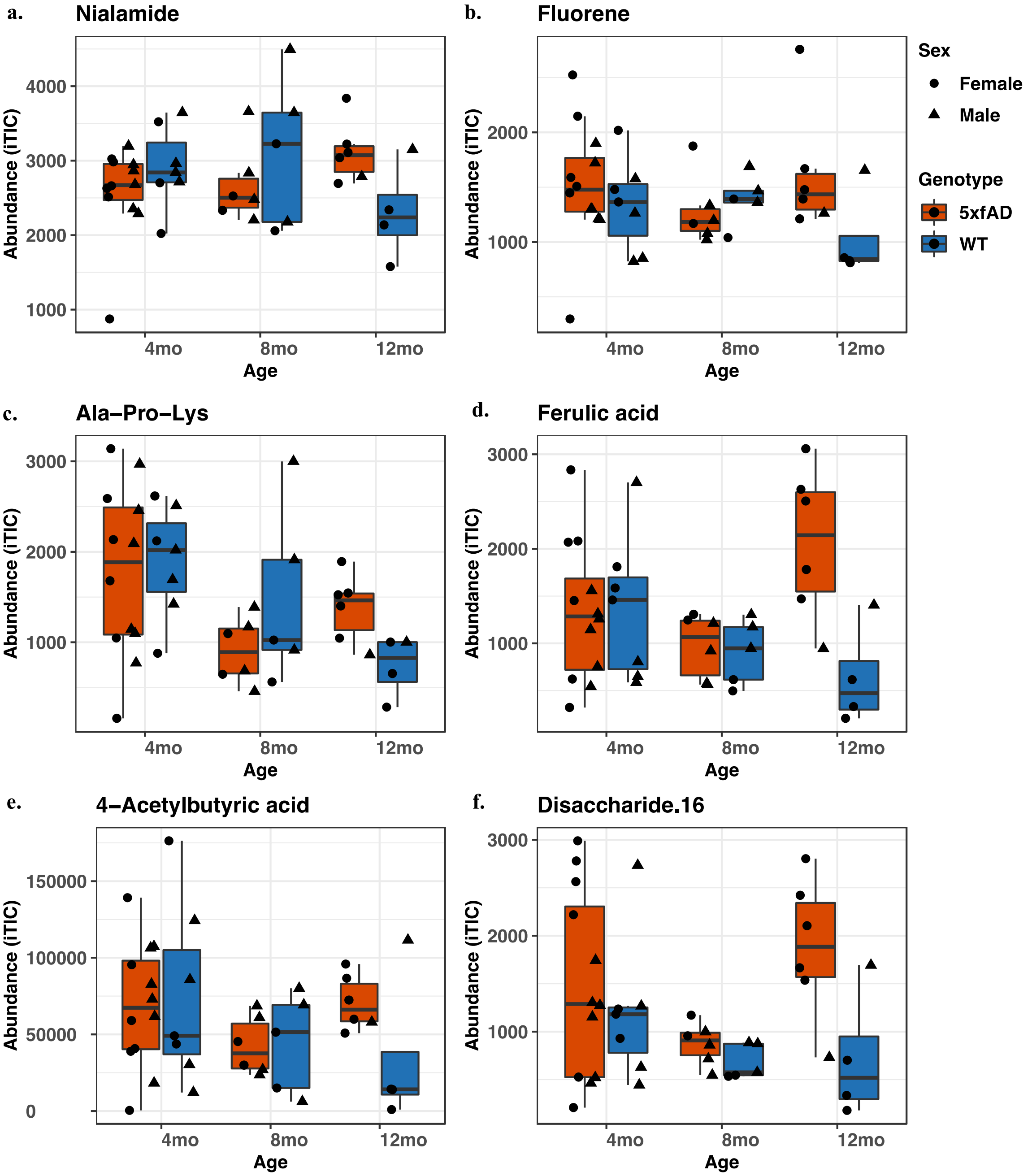


**Figure SR1**. Normalized abundance of the six fecal metabolites found to be significantly different at 12 months (females only) by LME (padj < 0.05). Significance was also confirmed by the Wilcoxon signed rank test ($p < 0.05$).

A correlation between the 100 most abundant metabolites and 17 significant species at 18 months of age (as revealed by the microbiome LME analysis shown in main text **Table 1**) shows distinct patterns of co-expressed microbes and bacteria, with a Spearman's co-occurrence range from –0.7 – 0.5 (**Figure SR2**). *Burkholderiales* bacterium YL45 was strongly anticorrelated with the first cluster of metabolites (N-omega-acetylhistamine, homovanillic acid sulfate, 2-(N-morpholino)ethanesulfonic acid, and 3-hydroxykynurenine) and co-occurred with the third (comprised of 23 metabolites), fifth (consisting only of 2'-deoxyadenosine), and 6$^{th}$ (containing 25 metabolites) clusters. *Ligilactobacillus murinus*, *Ligilactobacillus animalis*, and *Lactobacillus* sp. P38 were anticorrelated with the first and fourth metabolite clusters and co-occurred with the fifth cluster. The correlation patterns for *Lactobacillus johnsonii* stood apart from *Lactobacillus* sp. P38 and the two *Ligilactobacillus* species, showing anticorrelation with most metabolites except for the four molecules comprising the first cluster. *Romboutsia ilealis* and *Turicibacter* sp. H121 were weakly associated with most metabolites, and did not show pronounced correlation or anticorrelation patterns. The correlation patterns of the last two bacterial groupings are similar apart from the first metabolite cluster.

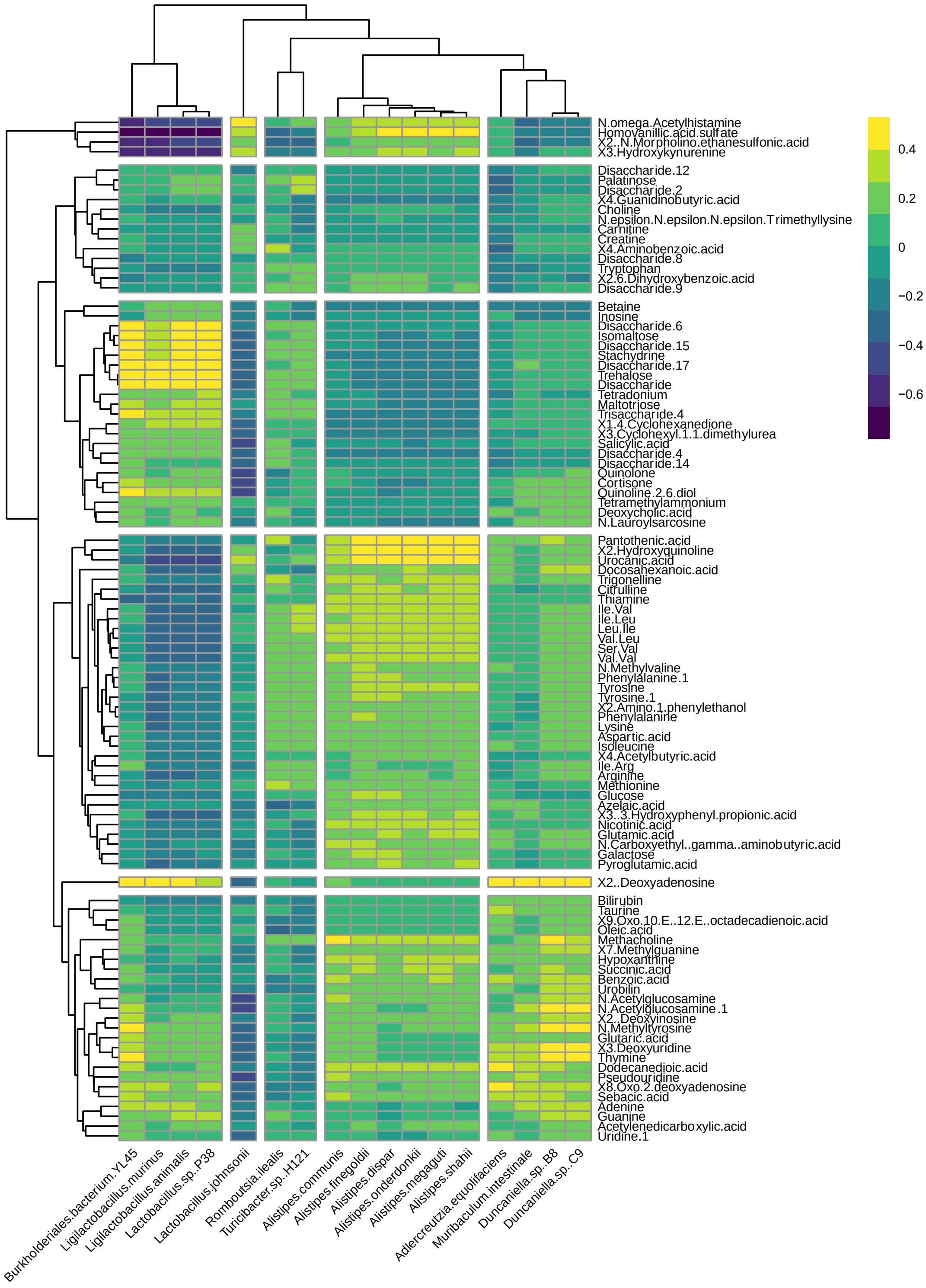


**Figure SR2**. Spearman correlation between fecal microbiome and metabolome (subset of samples from 4, 8, and 12 month cohorts).
